# BMAL1 loss in oligodendroglial lineage cells dysregulates myelination and sleep

**DOI:** 10.1101/2022.04.28.489946

**Authors:** Daniela Rojo, Anna Badner, Louisa Dal Cengio, Samuel Kim, Noriaki Sakai, Jacob Greene, Ella Eisinger, Caroline Arellano-Garcia, Lindsey C. Mehl, Mohammad E. Gumma, Rebecca L. Soyk, Julia Ransom, Maya K. Weigel, Belgin Yalçın, Samuel E. Jones, Hanna M. Ollila, Seiji Nishino, Erin M. Gibson

## Abstract

Myelination depends on maintenance of oligodendrocytes that arise from oligodendrocyte precursor cells (OPCs). We show that the dynamic nature of oligodendroglia and myelination are regulated by the circadian transcription factor BMAL1. *Bmal1* knockdown in OPCs during development – but not adulthood – decreases OPC proliferation, whereas BMAL1 regulates OPC morphology throughout life. OPC-specific *Bmal1* deficiency impairs remyelination in an age-dependent manner, suggesting that age-associated decrements in circadian regulation of oligodendroglia may contribute to the deficient remyelination potential in demyelinating diseases like multiple sclerosis (MS). This oligodendroglial dysregulation and dysmyelination increase sleep fragmentation in OPC-specific *Bmal1* knockout mice, and sleep fragmentation is causally associated with MS. These findings have broad mechanistic and therapeutic implications for numerous brain disorders that include both myelin and sleep phenotypes.

**One-Sentence Summary:** BMAL1 regulates the homeostatic maintenance of oligodendroglia and myelin, that subsequently controls sleep architecture.

## Main Text

Myelin ensheathes axons to facilitate efficient transduction of electrical signals and metabolic support of neurons (*1, 2*). Myelin-forming oligodendrocytes arise from oligodendrocyte precursor cells (OPCs), which are evenly distributed throughout the entire central nervous system (CNS). OPCs are highly dynamic as the most mitotic cells in the CNS (*3, 4*) with elaborate morphology that promotes motile filopodia and migration. OPC proliferation is stimulated when oligodendroglial loss initiates adjacent OPCs to divide (*5*), in response to neuronal activity in some neural circuits (*6*), and through biophysical and spatial constraints of their microenvironment (*7*). Even though OPCs are a functionally, spatially, and temporally heterogenous precursor population (*8–10*), they have a remarkable ability to maintain this consistent homeostatic density throughout the brain. How this unique harmony between prolific cellular self-renewal and population level homeostasis is achieved in normative brain health and disrupted in myelin-associated diseases like multiple sclerosis (MS) remains to be fully determined.

From cyanobacteria to humans, temporally dynamic mechanisms are imperative to the maintenance of homeostatic states and behaviors. This is afforded to organisms through the evolution of the circadian system which allows for biological processes to occur at the proper time of day. At the cellular level, circadian rhythms are generated ubiquitously throughout the body by a molecular transcriptional/translational feedback loop that has a period of ∼24 hours. Briefly, the products of the core clock genes—*Clock* and *Bmal1*—heterodimerize and drive the transcription of the clock genes families *Period* (*Per*) and *Cryptochrome* (*Cry*). Accumulated levels of PER and CRY within the cytoplasm feed back into the nucleus, displacing the CLOCK and BMAL1 heterodimer and consequently disrupting their own transcription (*11*). This transcriptional machinery regulates cytoskeletal factors (*12*), cell cycle (*13*), and metabolism (*14*) in numerous cell populations. OPCs proliferate on a circadian cycle (*4*) and sleep deprivation negatively impacts OPC proliferation and differentiation (*15*). The potential role this circadian clock plays in regulating the homeodynamic nature of oligodendroglial lineage cells remains unknown. We posit that the molecular circadian system driven by the transcription factor BMAL1 regulates oligodendroglial cells and myelination, which contribute to the maintenance of systems-level homeostatic processes such as sleep.

## Results

### Functional knock down of circadian gene Bmal1 dysregulates OPC proliferation and morphology

To determine the role of the BMAL1-driven molecular clock in regulating OPC biology, we first confirmed expression of the complete molecular clock machinery throughout the oligodendroglial lineage using the RNAseq transcriptome database of cell type-specific gene expression in mouse brain (*16, 17*) (**Fig. S1**), and we confirmed BMAL1 expression in mouse OPCs (**Fig. 1A-B**). We next developed a conditional clock gene knockout (*Bmal1*^*fl/fl*^) and cell type-specific Cre driver mouse model (*NG2::Cre*) to constitutively eliminate functional *Bmal1* from OPCs during embryonic development (**Fig. 1C**). BMAL1 levels were significantly decreased in OPCs isolated from postnatal day (P)6 *NG2::Cre+*;*Bmal1*^*fl/fl*^ (OPC-*Bmal1*-KO) compared to *NG2::Cre-*;*Bmal1*^*fl/fl*^ (OPC-*Bmal1*-WT) mice (**Fig. 1D**). We hypothesized that BMAL1 loss in OPCs would disrupt OPC dynamics given that it regulates cell cycle, proliferation and cytoskeletal factors in other cells (*12, 13*). Mice were injected with the thymidine analogue 5-ethynyl-2’-deoxyuridine (EdU) on P18-20 to mark newly proliferated OPCs. EdU^+^/PDGFRα^+^ OPCs were significantly reduced in the corpus callosum of P21 OPC-*Bmal1*-KO compared to OPC-*Bmal1*-WT mice (**Fig. 1E-F**). *Bmal1* loss led to a significant decrease in OPC density of OPC-*Bmal1*-KO compared to OPC-*Bmal1*-WT mice (**Fig. 1G-H**) that persisted into adulthood (P63; 5527 ± 221.7 vs 4289 ± 355 cells/mm^3^; *n* = 4-6 mice; *P* < 0.05). These cellular differences did not result in corpus callosum volume changes (**Fig. S2A**). We also identified a significant reduction in OPC morphological complexity by evaluating the length, volume, and branching points of processes in OPC-*Bmal1*-KO compared to OPC-*Bmal1*-WT mice (**Fig. 1I-J**).

**Fig. 1.**
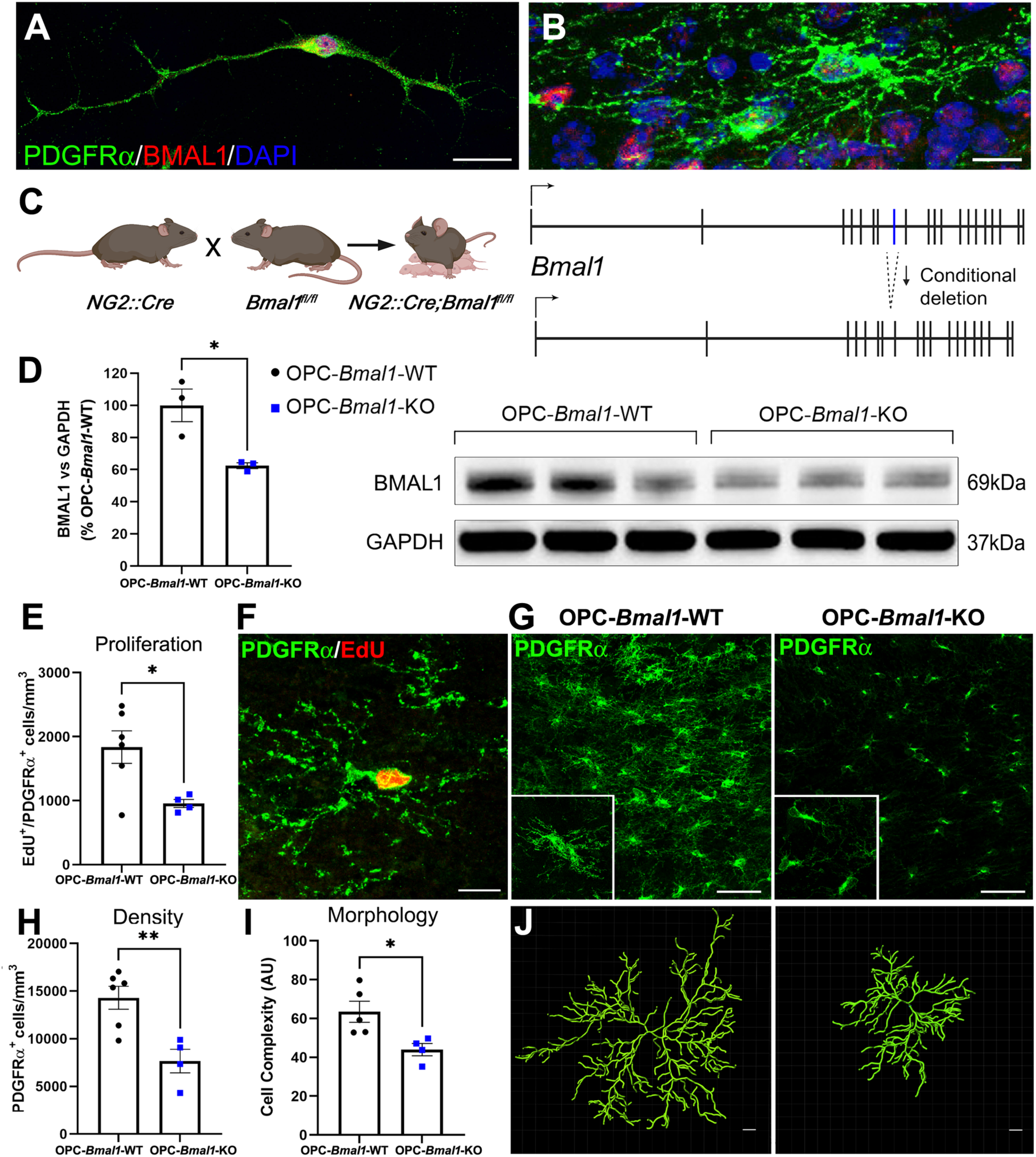
Knock out of *Bmal1* in OPCs affects proliferation and morphology. **(A-B)** Photomicrograph (63X) of PDGFRα^+^ OPCs (green) expressing BMAL1 (red) with DAPI (blue) in immunopan-isolated OPCs from P6 mice; Scale bar = 20µm **(A)**, and in the mouse brain; Scale bar = 10µm **(B). (C)** The conditional *Bmal1* knock out model *Bmal1*^*fl/fl*^, in which the eighth exon coding for the basic helix-loop-helix DNA-binding domain is surrounded by loxP sites, was bred to the OPC-specific *Cre* driver mouse model *NG2::Cre* to constitutively knock out functional BMAL1 binding in OPCs (*NG2::Cre*+*;Bmal1*^*fl/fl*^ or OPC-*Bmal1-*KO). **(D)** Western blot of immunopan-isolated OPCs from P6 mice. OPCs lacking *Bmal1* express 38% less BMAL1 than *Bmal1* intact (*NG2::Cre*-*;Bmal1*^*fl/fl*^ or OPC-*Bmal1-*WT) OPCs (n = 3 per group). **(E-F)** Loss of *Bmal1* from OPCs results in a 48% decrease in OPC proliferation assessed by EdU^+^/PDGFRα^+^ co-labelling in the corpus callosum at P21 **(E). (F)** Photomicrograph (63X) of PDGFRα^+^ (green) OPC with EdU (red). Scale bar = 10µm. **(G)** Photomicrographs (20X; inset 63X) of PDGFRα^+^ (green) OPCs in the corpus callosum at P21. Scale bar = 50µm. **(H)** OPC-*Bmal1-*KO mice exhibit a 46% decrease in PDGFRα^+^ OPC density **(G)** compared to OPC-*Bmal1-*WT OPCs in the corpus callosum (n = 4-6 per group). **(I)** Morphological complexity (ratio of branch points and filament length by filament volume) of OPCs is decreased by the lack of *Bmal1* (n = 4-5 per group). **(J)** Imaris analysis representation. OPC-*Bmal1-*WT PDGFRα^+^ OPCs maintain proper density and arborization (left) compared to reduced complexity in OPC-*Bmal1-*KO OPCs (right). Scale bar = 5µm. Data shown as mean +/- SEM. * p<0.05, ** p<0.01.

### OPC-specific Bmal1 knock down dysregulates oligodendrocytes and myelination

Given the striking effects of BMAL1 loss on OPCs, we next tested how loss of *Bmal1* affects myelinating oligodendrocytes. Because a decrease in OPC density impacts the progenitor pool available for oligodendrogenesis and considering oligodendrocytes also lack functional BMAL1 in our model due to embryonic knock down, we evaluated the effect of OPC-specific *Bmal1* loss on oligodendrogenesis. OPC-*Bmal1*-KO mice exhibit a significant decrease in density of CC1^+^ oligodendrocytes and myelin basic protein (MBP) content in the corpus callosum at P21, an age that corresponds to the end of developmental myelination (**Fig. 2A-C**). Using transmission electron microscopy (TEM), we observed thinner myelin sheaths quantified as an increase in the *g*-ratio (*g*-ratio = inner axon diameter/total fiber diameter) of axons projecting from the cortex into the corpus callosum of the premotor circuit in OPC-*Bmal1*-KO compared to control mice (**Fig. 2D-F, Fig. S3**). This decrease in myelin sheath thickness was found in small and medium but not large caliber axons (**Fig. 2E**). As NG2 is a proteoglycan that is not only expressed in OPCs but also in pericytes, *Bmal1* loss in NG2^+^ cells could potentially compromise the blood-brain barrier (BBB), leading to brain inflammation and neurotoxicity promoting oligodendroglia and myelin loss. Brain-wide loss of *Bmal1* has previously been linked to BBB hyperpermeability associated with pericyte dysfunction and dysregulation of platelet-derived growth factor receptor β (*18*). To rule out this possibility, we tested the integrity of the BBB through administration of sodium fluoresceine and found no differences in brain permeability between genotypes at P21 (**Fig. S2B**).

**Fig. 2.**
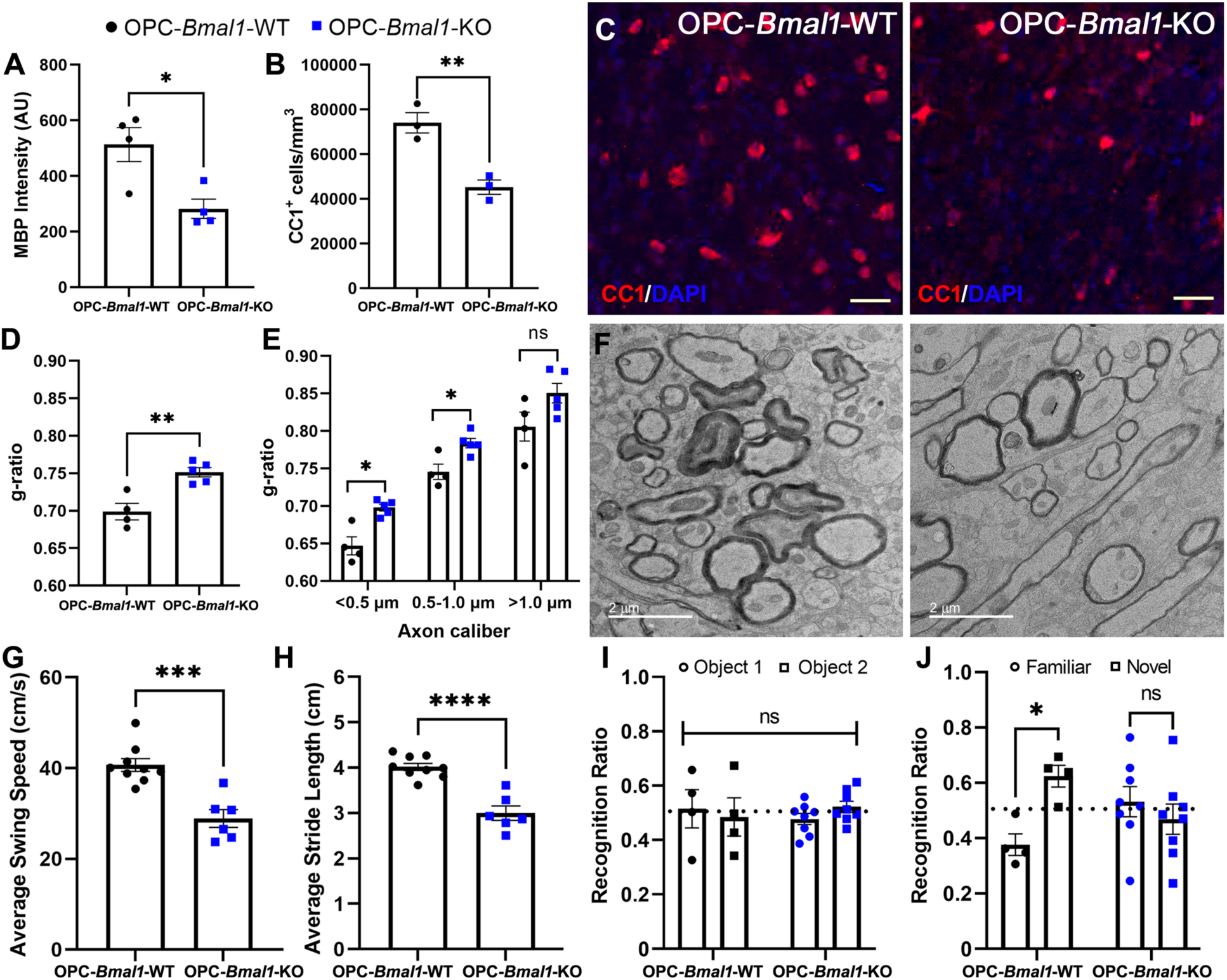
Constitutive knock out of *Bmal1* in OPCs leads to deficits in oligodendrocytes, myelination, and motor and cognitive function. **(A-C)** Constitutive knock out of *Bmal1* in OPCs (OPC-*Bmal1-*KO) results in decreased myelin basic protein (MBP) content measured by immunofluorescent intensity **(A)** and CC1^+^ oligodendrocyte density **(B)** in the corpus callosum at P21 compared to OPC-*Bmal1-*WT (n = 3-4 per group). **(C)** Photomicrographs (20X) of CC1^+^ (red) oligodendrocytes with DAPI (blue). Scale bar = 30 µm. **(D)** These reductions are associated with a decrease in myelin sheath thickness at P21 (larger g-ratio = thinner myelin). **(E)** g-ratio by axon caliber (n = 4-5 per group). **(F)** Representative TEM images of cortical projections to corpus callosum in OPC-*Bmal1-*WT (left) and OPC-*Bmal1-*KO (right) mice. Scale bar = 2 µm. **(G-H)** OPC-*Bmal1-*KO exhibit deficits in swing speed of limbs **(G)** and shorter stride length **(H)** at P35 assessed using the CatWalk gait analysis system compared to OPC-*Bmal1-*WT mice (n= 6-9 group). **(I-J)** OPC-*Bmal1-*KO mice exhibit attention and short-term memory deficits at 7 months assessed using a modified novel object recognition test (NORT) in which the interval between training **(I)** and testing **(J)** is shortened to 5 minutes (n = 4-8 group). **(J)** OPC-*Bmal1-*KO mice do not discriminate between the novel and familiar objects whereas OPC-*Bmal1-*WT spend more time investigating the novel over familiar object. Data shown as mean +/- SEM. ns p>0.05, * p<0.05, ** p<0.01, ***p<0.001, **** p<0.0001.

### Loss of Bmal1 in oligodendroglial lineage cells disrupts motor and cognitive function

We next evaluated if these cellular and myelin decrements contributed to alterations in motor and cognitive behaviors known to be affected by changes in frontal lobe white matter (*19, 20*). The identified decrements in OPC-*Bmal1*-KO mice produced a significant dysregulation of motor behavior at P35, including a decrease in paw swing speed and stride length assessed using the CatWalk gait analysis system (**Fig. 2G-H**). We also tested cognition using a modified novel object recognition test (NORT) that places greater emphasis on attention and short-term memory instead of hippocampal-dependent long-term memory (*19, 20*). OPC-*Bmal1*-WT mice spent more time investigating the novel over familiar object while OPC-*Bmal1*-KO mice did not discriminate between the objects during the testing phase, suggesting deficits in white matter-associated cognition (**Fig. 2I-J**).

### BMAL1-mediated regulation of OPCs proliferation and differentiation in vitro

To understand the molecular mechanism(s) through which BMAL1 controls OPCs, we first confirmed rhythmic clock gene expression in OPCs isolated from P6 *Per2-Luciferase* mice that express luciferin under the control of *Per2*. We synchronized OPCs *in vitro* using 100 nM dexamethasone and confirmed circadian synchronization by measuring luminescence every 4 hrs (**Fig. 3A**). The expression rhythmicity of the clock genes *Bmal1, Per2*, and *Rev-Erbα* (a transcriptional repressor of *Bmal1*) was dampened in OPCs isolated from OPC-*Bmal1*-KO (**Fig. 3B**) compared to rhythmic expression in OPC-*Bmal1*-WT mice, confirming global circadian dysregulation of clock gene expression in *Bmal1* knockout OPCs. We then interrogated if the cellular effects of *Bmal1* loss in OPCs were a consequence of decreased OPC number exclusively or potentiated by a dysregulation of their differentiation potential. In OPCs isolated from OPC-*Bmal1*-KO mice, we found a 14% decrease in proliferating EdU^+^/PDGFR*α*^+^ cells in *Bmal1* knockout compared to control OPCs (**Fig. 3C-D**). We then induced OPC differentiation for 3 days and found a decrease in MBP^+^ oligodendrocytes when *Bmal1* was knocked down (**Fig. 3E-F**) that recovered when differentiation was completed after 6 days in both genotypes (**Fig. 3G-H**).

**Fig. 3.**
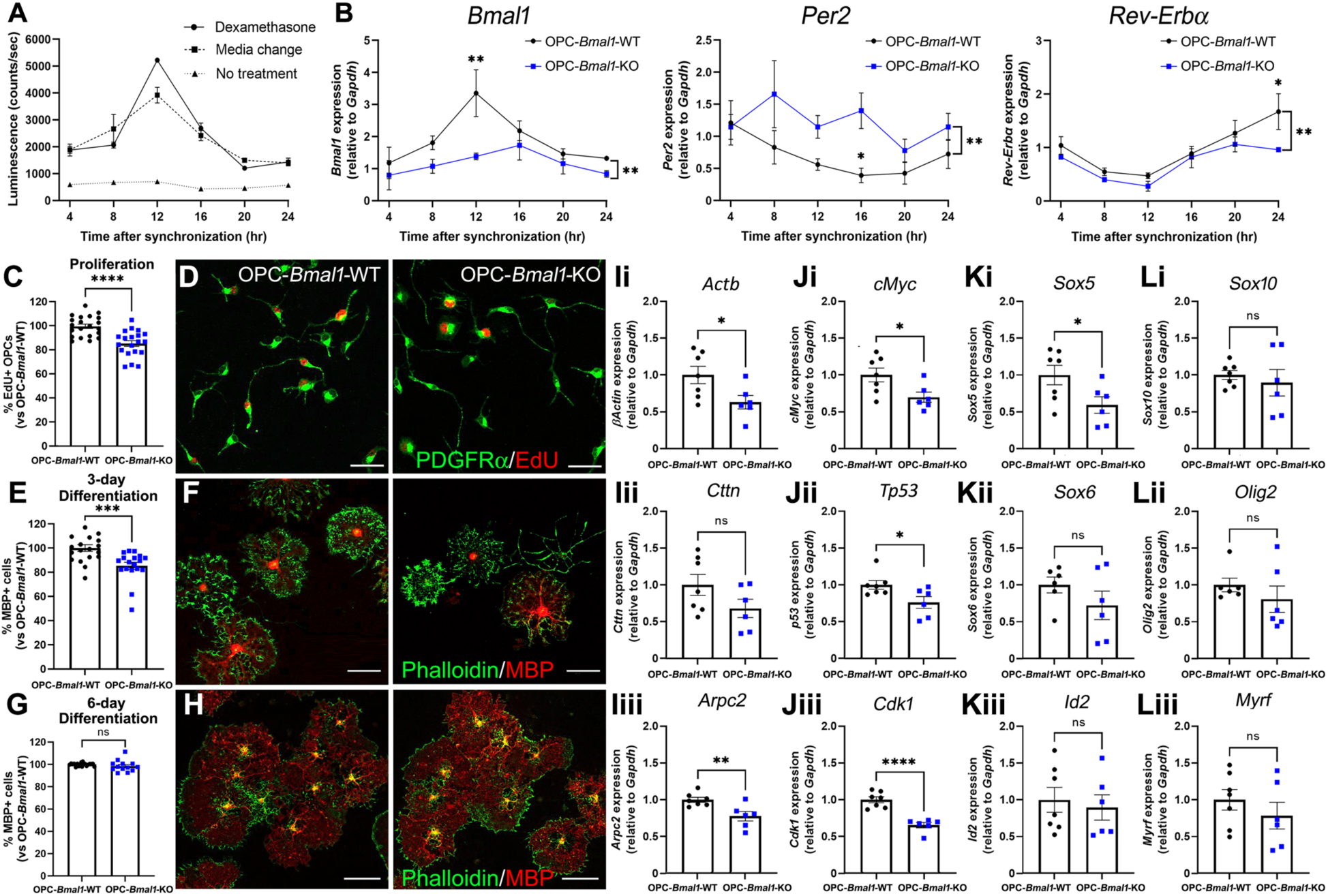
BMAL1 regulates genes associated with OPC cycling, proliferation and morphology but not complete differentiation. **(A)** Luciferase activity under the control of *Per2* promoter in isolated OPCs from P6 *Per2::Luciferase* mice. Treating OPCs with 100 mM dexamethasone for 1 hr or implementing a media change containing 10 µM forskolin synchronizes the circadian molecular clocks of OPCs *in vitro* (n = 4 per group). **(B-C)** Cycling of clock genes in OPCs synchronized *in vitro* is inhibited by knocking out *Bmal1* (OPC-*Bmal1-*KO). **(B)** *Bmal1, Per2*, and *Rev-Erbα* expression measured by RT-qPCR relative to *Gapdh* across 24 hr circadian time (CT) (n = 4 per group). **(C)** Knock out of *Bmal1* leads to a 14% decrease in proliferation as marked by incorporation of EdU compared to circadian intact OPCs (OPC-*Bmal1-*WT) (N=4 experimental replicates with n=3-8 per genotype). **(D)** Photomicrographs (40X) of PDGFRα^+^ (green) OPCs incubated with EdU (red) for 5 hrs. Scale bar = 40 µm. **(E-H)** OPC-*Bmal1-*KO mice exhibit a delay in OPC differentiation after 3 days of induction **(E)** that recovers after 6 days of differentiation **(G)** (N=3 experimental replicates with n=3-8 per genotype). **(F, H)** Photomicrographs (20X) of oligodendroglia lineage cells after 3 days **(F)** or 6 days **(H)** of differentiation induction showing MBP expression (red) and actin filament staining with phalloidin (green) to distinguish the differentiation stage. Scale bar = 50 µm **(F)** and 100 µm **(H). (I)** OPC-*Bmal1-*KO mice express less *Actb* and *Arpc2* cytoskeleton genes, measured by RT-qPCR in immunopan-isolated OPCs at zeitgeber time (ZT)6 (ZT0=lights on; ZT12=lights off). **(J-L)** The decrease in OPC proliferation identified in OPC-*Bmal1*-KO mice is associated with a decrease in expression of genes that control cell cycle **(J)** and OPC proliferation **(K)**, but not differentiation **(L)** (n = 6-7 per genotype). Data shown as mean +/- SEM. ns p>0.05, *p<0.05, ** p<0.01, *** p<0.001, ****p<0.0001.

As the *Bmal1*-driven circadian clock can regulate from 10-50% of the genome (*21*), it is possible that the effects observed in the OPC-specific *Bmal1* knockout are due to a variety of downstream pathways controlled by BMAL1. BMAL1-associated OPC deficits are primarily related to decreased expression in genes linked with cytoskeletal regulation (*Actb* and *Arpc2*; **Fig. 3I**,) cell cycle (*cMyc, Tp53* and *Cdk1*; **Fig. 3J**), and proliferation (*Sox5*; **Fig. 3K**), but not differentiation (*Sox10, Olig2* and *Id2*; **Fig. 3L**).

### Effect of BMAL1 disruption in OPCs during adulthood

As the two major waves of developmental OPC migration that begin OPC density establishment occur during mid-embryonic development (*22*) and developmental oligodendrogenesis is completed before early adulthood in mice, we hypothesized that *Bmal1* disruption in OPCs in adulthood would result in less dramatic effects on OPC dynamics. To investigate this, we induced knock out of *Bmal1* in OPCs using a conditional, inducible OPC-specific Cre mouse line *PDGFR*α*::CreER*^*T2*^ crossed with *Bmal1*^*fl/fl*^. *PDGFR*α*::CreER*^*T2*^+;*Bmal1*^*fl/fl*^ (OPC-*Bmal1*-iKO) and *PDGFR*α*::CreER*^*T2*^-;*Bmal1*^*fl/fl*^ (OPC-*Bmal1*-WT) mice were injected intraperitoneally with 100 mg/kg tamoxifen for 3 consecutive days at 3 months (Younger Adults) or 10 months (Older Adults) of age and brains were assessed 6 weeks later. We obtain approximately 80% recombination following this 3 day tamoxifen schedule (*19*), and found no evidence of ‘leaky’ Cre expression in neurons in this Cre driver (*23*). We found that OPC density is no longer decreased, but morphological complexity is still disrupted when *Bmal1* knock out is induced in younger and older adults (**Fig. 4A-B**) rather than embryonically as above. Taken together, our findings illustrate a key role for *Bmal1* in oligodendroglial lineage homeostasis and function during development and in adulthood.

**Fig. 4.**
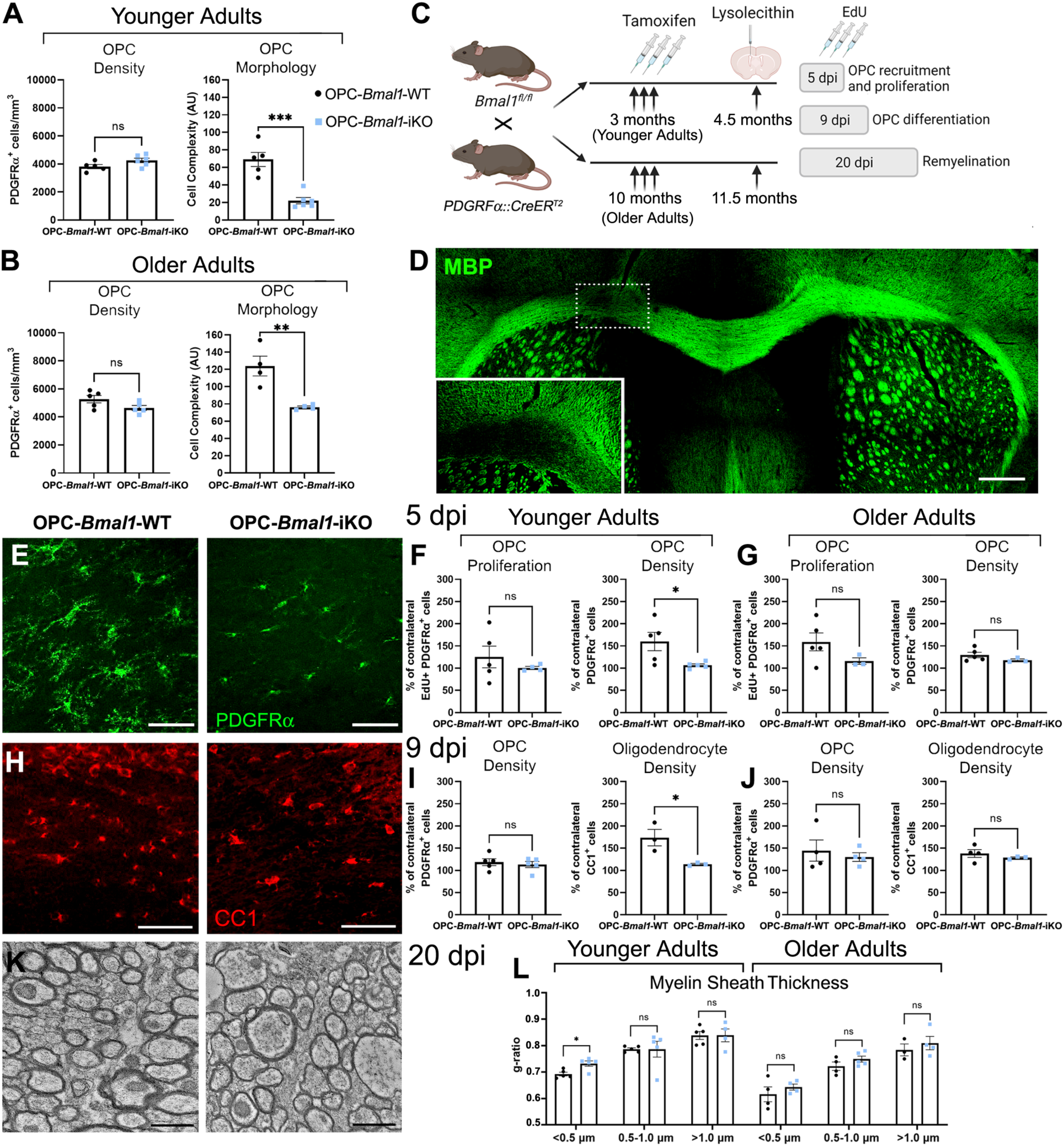
BMAL1 disruption during adulthood impairs OPC response to demyelination. **(A-B)** Tamoxifen-induced knock out of *Bmal1* in OPCs (*PDGFRα::CreER*^*T2*^*+;Bmal1*^*fl/fl*^ or OPC-*Bmal1-* iKO) at 3 (Younger Adults) **(A)** or 10 (Older Adults) months of age **(B)** leads to decreased complexity in OPC morphology without disrupting OPC density. **(C)** 6 weeks after tamoxifen-induced Cre recombination, OPC-*Bmal1-*iKO and their control littermates (*PDGFRα::CreER*^*T2*^*-;Bmal1*^*fl/fl*^ or OPC-*Bmal1-*WT) were injected with lysolecithin into the cingulum of the corpus callosum of one hemisphere and brains were collected after 5, 9 or 20 days post-injection (dpi). Cellular density and remyelination were compared to the contralateral non-lesioned hemisphere. **(D)** Photomicrograph (10X) of MBP (green) showing the demyelinating lesion in the cingulum of the corpus callosum. Scale bar = 500µm. **(E)** Photomicrographs (20X) of PDGFRα^+^ (green) OPCs in the corpus callosum of younger adults at 5dpi. Scale bar = 50µm. **(F)** 5 days after demyelination, OPC density in the lesion compared to the contralateral non-lesioned hemisphere is lower in younger adult OPC-*Bmal1-*iKO than OPC-*Bmal1-*WT mice, which is unrelated to an increase in OPC proliferation. **(G)** No significant differences in OPC density are observed between OPC-*Bmal1-*WT and OPC-*Bmal1-*iKO older adults at 5 dpi. **(H)** Photomicrographs (20X) of CC1^+^ (red) oligodendrocytes in the corpus callosum of younger adults at 9 dpi. Scale bar = 50µm. **(I-J)** 9 days after lysolecithin-induced demyelination, oligodendrocyte density in the lesion is significantly lower in OPC-*Bmal1-*iKO compared to OPC-*Bmal1-*WT mice in younger **(I)**, but not in older adults **(J)** with not differences in OPC proliferation at either age. **(K)** Representative TEM images of the corpus callosum in OPC-*Bmal1-*WT and OPC-*Bmal1-*iKO younger adults at 20 dpi. Scale bar = 1 µm. **(L)** At 20 dpi, myelin sheath thickness is decreased in low caliber axons of OPC-*Bmal1-*iKO compared to OPC-*Bmal1-*WT younger adults. Data shown as mean +/- SEM. ns p>0.05, * p<0.05, ** p<0.01, *** p<0.001.

### Remyelination potential of oligodendroglial lineage cells lacking Bmal1

Deficits in OPC density and differentiation are underlying causes for limited remyelination in MS (*24*). Given that *Bmal1* loss in OPCs results in pronounced cellular changes during developmental myelination, we hypothesized that the OPC circadian clock is necessary for proper cellular recovery following a demyelinating injury. We generated a unilateral focal demyelinating lesion in the cingulum of the corpus callosum with stereotactic injection of lysolecithin 6 weeks after tamoxifen injections at 3 or 10 months of age in OPC-*Bmal1*-iKO and OPC-*Bmal1*-WT mice (**Fig. 4C-D**). Lysolecithin-induced lesions progress from an active demyelination phase during the first 3 days, to OPC recruitment (days 3 to 7), differentiation (days 7 to 10), and active remyelination (days 10 to 21) (*25*). We first evaluated OPC proliferation and density in the lesion 5 days post-lysolecithin injection (dpi). When *Bmal1* knockout and demyelination occur in young adulthood, intra-lesion OPC density is significantly lower in OPC-*Bmal1*-iKO mice than OPC-*Bmal1*-WT mice. This difference was not attributed to OPC proliferation changes (**Fig. 4E-F**). By 9 dpi the OPC density difference no longer exists between genotypes, but the number of lesion-associated oligodendrocytes compared to the non-lesioned hemisphere is significantly decreased in OPC-*Bmal1*-iKO compared to OPC-*Bmal1*-WT young adults (**Fig. 4H-I**). Twenty days after demyelination, remyelination of small caliber axons is reduced in OPC-*Bmal1*-iKO compared to OPC-*Bmal1*-WT (**Fig. 4K-L**). When *Bmal1* loss and demyelination are induced in older adults, the significant differences between genotypes in OPCs at 5 dpi (**Fig. 4G**), oligodendrocytes at 9 dpi (**Fig. 4J**) and remyelination at 20 dpi (**Fig. 4L**) were not found, suggesting that the ability of OPCs to dynamically respond to a demyelinating lesion is impacted by both age and BMAL1 status.

### Disruption of BMAL1 in oligodendroglia is associated with sleep fragmentation in mice

As the global elimination of *Bmal1* markedly disrupts circadian and sleep processes in mice (*26, 27*) and given the robust effects of even small changes in myelination on brain-wide neural circuit dynamics (*28*), we tested the hypothesis that *Bmal1* disruption of the oligodendroglial lineage may lead to systems-level dysregulation of homeostatic circuits regulating circadian and sleep processes. We thus assessed how constitutive *Bmal1* loss in OPCs affects global circadian rhythmicity and sleep during adulthood. We subjected OPC-*Bmal1*-WT and OPC-*Bmal1*-KO mice to a 12::12 light/dark (LD) cycle for 7 days to assess circadian entrainment followed by constant darkness (DD) for 15 days to assess free-running circadian rhythms (**Fig. 5A**). OPC-*Bmal1*-KO mice exhibit normal circadian periods (tau) compared to controls (**Fig. S4A**) with no genotype effect on locomotor activity throughout the day (**Fig. S4B-C**). At 3.5 months of age, we implanted electroencephalogram (EEG) electrodes into the cortex of OPC-*Bmal1*-WT and OPC-*Bmal1*-KO mice to measure sleep waves across the hippocampus and evaluated baseline sleep recordings in light::dark cycles. In mice, sleep is polyphasic, and around two thirds of these short sleep periods occur during the light/rest cycle. Sleep is characterized by two main stages consisting of rapid eye movement (REM) and non-REM (NREM) sleep. The general pattern of sleep/wake is not affected in mice with *Bmal1*-disrupted OPCs, as the total time spent awake does not vary between genotypes during the light or dark phases (**Fig. 5B left**). However, when we evaluate sleep architecture, *Bmal1* loss in OPCs leads to significant sleep fragmentation during the dark/active phase; OPC-*Bmal1*-KO mice have 29% shorter but 57% more frequent wake events (**Fig. 5C-D left, H**). During the dark/active phase, NREM events are also shorter but more frequent even though the total time spent in NREM does not differ (**Fig. 5E-G left, H**). REM sleep does not vary between genotypes (**Fig. S5A-C**). To further elucidate the effect of OPC-specific BMAL1 on sleep homeostasis, we subjected the mice to a 6-hr sleep deprivation challenge. The effect on sleep fragmentation is exacerbated in OPC-*Bmal1*-KO mice following sleep deprivation as wake events during the dark phase are 49% shorter and 95% more frequent than controls (**Fig. 5B-D right**). The total time spent in NREM during the dark/active phase is significantly increased in OPC-*Bmal1*-KO compared to OPC-*Bmal1*-WT mice and the events are shorter and more frequent, following the same pattern as baseline (**Fig. 5E-G right**). Taken together, these findings show that the genetic disruption of *Bmal1* in oligodendroglial lineage cells leads to increased sleep fragmentation and a greater drive for restorative NREM sleep following sleep deprivation, illustrating a role for oligodendroglia in sleep function.

**Fig. 5.**
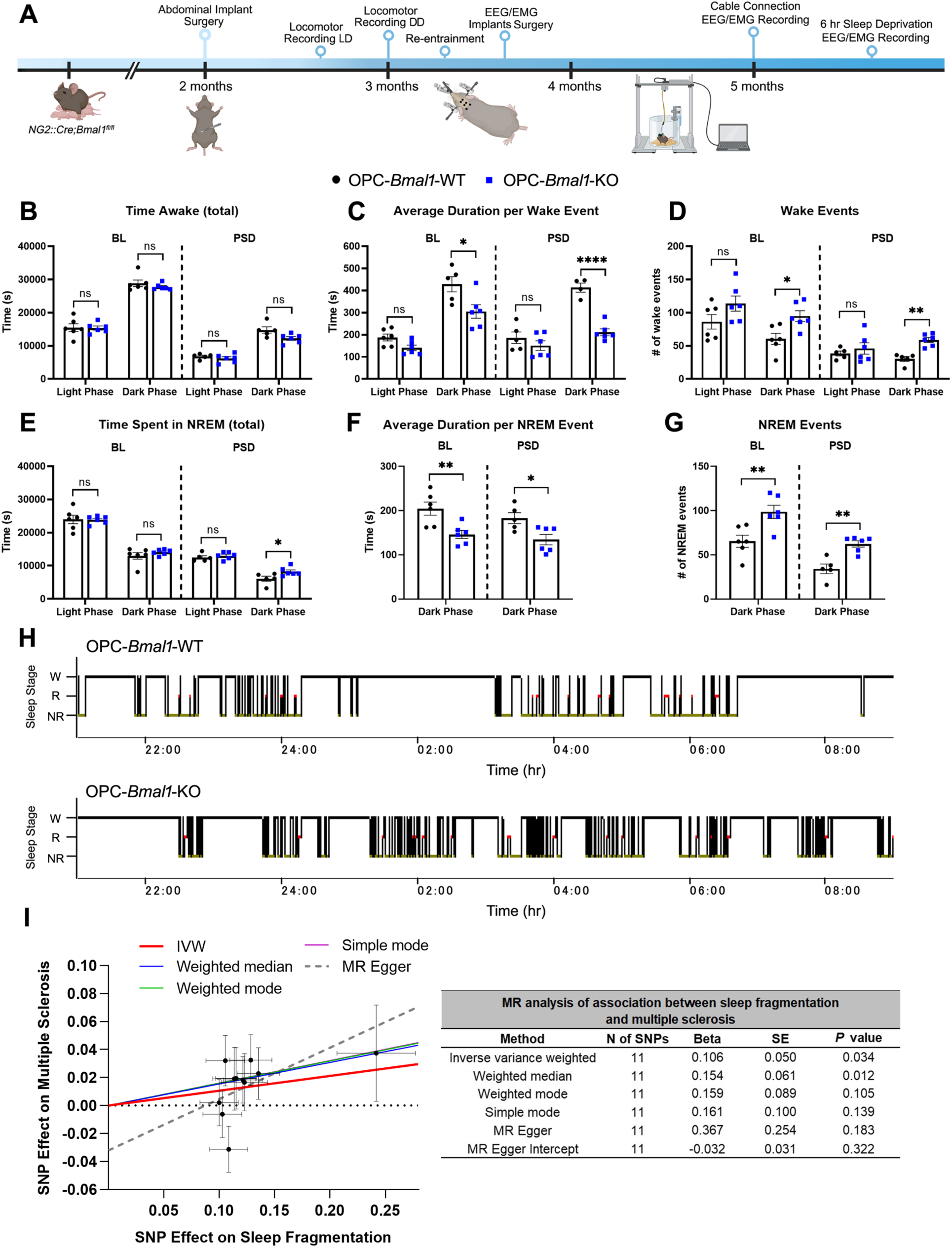
Sleep fragmentation is associated with aberrant myelination in mice and individuals with multiple sclerosis. **(A)** At 2 months of age, OPC-*Bmal1-*WT and OPC-*Bmal1-*KO mice were subjected to abdominal surgery to implant electronic transmitters. Locomotor activity was telemetrically monitored in light/dark (LD) cycles for 7 days followed by constant darkness (DD) for 15 days. EMG/EEG biotelemetry electrodes were then implanted at 3.5 months and baseline (BL) sleep was recorded at 5 months for 10 days. Mice were then subjected to a 6-hr sleep deprivation (SD) cycle and sleep waves were recorded for 18 hrs post sleep deprivation (PSD). **(B-D)** While OPC-*Bmal1-*WT and OPC-*Bmal1-*KO mice spend the same total amount of time awake and asleep during both the light/sleep and dark/active phase at baseline **(B)**, OPC-*Bmal1-*KO mice have shorter **(C)**, more frequent wake bouts **(D)** during the dark phase than OPC-*Bmal1-*WT mice, indicative of sleep fragmentation. Following SD, OPC-*Bmal1-*KO mice exhibit a significant decrease in the amount of wake during their dark/active phase compared to OPC-*Bmal1-*WT mice **(C)**. This increased active-phase sleep post-sleep deprivation exhibits the same pattern of sleep fragmentation as detected during baseline (i.e. shorter but more frequent bouts). **(E-G)** The shift in sleep architecture at BL is driven by changes in NREM sleep, as total amount of NREM does not differ between the genotypes **(E)**, but OPC-*Bmal1-*KO mice exhibit the same trend of shorter **(F)** but more frequent **(G)** NREM events during the dark/active phase. Following SD, OPC-*Bmal1-* KO spend more time in NREM sleep compared to OPC-*Bmal1-*WT mice **(E)**, which is driven by shorter **(F)**, more frequent **(G)** bouts. **(H)** Representative hypnograms showing wake (W), REM (R) and NREM (NR) sleep events during the dark phase (21:00 hr – 9:00 hr) at baseline in OPC-*Bmal1-*WT (top) and OPC-*Bmal1-*KO (bottom) mice. n= 4-6 per group. **(I)** Scatterplot of the mendelian randomization (MR) analysis of association between sleep fragmentation and multiple sclerosis (MS). The y-axis represents the effect of the analyzed variants on MS (beta) and the x-axis represents the variants effect on the number of sleep episodes (logOR) for each of the 11 variants studied. The slopes of the regression lines represent the causal association tested using Inverse variance weighted (IVW), Weighted median, Weighted mode, Simple mode, and MR Egger statistical tests. Corresponding table of the MR analysis performed indicates causal association between sleep fragmentation and MS risk without pleiotropic effect. Beta, β coefficient; SE, Standard Error. Data shown as mean +/- S.E.M. n.s p>0.05, *p<0.05, **p<0.01, ****p<0.0001.

### Association of sleep fragmentation with multiple sclerosis in humans

Given that dysmyelination in OPC-*Bmal1*-KO mice is associated with increased sleep fragmentation, could changes in sleep fragmentation be associated with myelin disorders in humans? We evaluated the causal association of changes in sleep on the risk of MS. We performed Mendelian randomization (MR) analyses on lead variants identified in a GWAS of sleep fragmentation (defined as number of sleep episodes) in 85,723 UK Biobank individuals (*29*). We evaluated their association with MS risk by leveraging results from a GWAS of MS in the International Multiple Sclerosis Genetics Consortium (N = 47,429 MS patients and 68,374 controls) (**Table S1**) (*30*). Eleven variants associated with number of sleep episodes/sleep fragmentation were suitable for MR analysis after exclusions (**Table S1**). We identified that sleep fragmentation is causally associated with an increased risk of MS (Inverse Variance Weighted OR = 1.11 [1.01-1.23], P = 0.034, per unit increase in the number of sleep episodes, 95%; **Fig. 5I**). Analysis using statistically weaker methods provided consistent causal and statistically significant effect at an α of 0.05 with Weighted median method (OR = 1.17 [CI = 1.03-1.32], P = 0.012, **Fig. 5I**). Furthermore, MR Egger analysis, which estimates the total pleiotropic effect of the instruments used, had a causal effect size with consistent direction of effect to the other four methods indicating that there was no significant pleiotropy detected (MR Egger Intercept P = 0.32; **Fig. 5I**). Further analysis using all variants without correcting for winner’s curse provided consistent causal and significant effect at an α of 0.05 with Weighted median method (OR = 1.14 [CI = 1.03-1.27], P = 0.016, **Fig. S6**). Taken together, these findings indicate a previously underappreciated association between sleep fragmentation and myelin disease.

## Discussion

The results presented here demonstrate a critical role for BMAL1 in modulating OPC dynamics and myelination throughout development, adulthood, and in disease. We show that BMAL1 is not only necessary for the dynamic nature of oligodendroglial lineage cells and myelination but also for the homeostatic regulation of sleep architecture. By eliminating *Bmal1* from OPCs, we find a significant deficit in density, proliferation, and morphological complexity, leading to a decrease in oligodendrocytes. These cellular deficits translate into thinner myelin sheaths and decrements in motor and cognitive functions associated with white matter structures (*20*). Importantly, our results suggest this oligodendroglial decrease is related to precursor population depletion more than altered differentiation, exemplified by diminished gene expression of cell cycle and proliferation regulators. This observation is in agreement with the known role of BMAL1 in regulation of cell cycle checkpoints in other cells (*13*). Previous work identified increases in transcripts that control OPC proliferation during the sleep/light phase (*15*), a period when *Bmal1* peaks in mice. As OPCs are the most consistently mitotic cells in the CNS (*3, 4*), even small changes in proliferation can disrupt their homeostatic density. The oligodendrocytes that arise from these OPCs will also putatively lack functional BMAL1 because of its constitutive loss in OPCs during embryonic development. While our *in vitro* studies indicate that BMAL1 regulates OPC proliferation and division more so than differentiation, future studies specifically targeting knock down of *Bmal1* in oligodendrocytes will further stratify the role of BMAL1 in myelination.

Even though BMAL1 continues to regulate OPC morphology throughout life, the effect on OPC density is abrogated when BMAL1 elimination is initiated during adulthood. As BMAL1 regulates the OPC cell cycle, and the mitotic rate of OPCs decreases with age (*9*), this finding further supports the role of BMAL1 in controlling OPC division. OPCs are morphologically dynamic with surveillance of their microenvironment. This allows them to not only establish and maintain their equal distribution but also to respond to loss of adjacent OPCs (*5*). The decrease in *Actb* expression in OPCs following BMAL1 loss suggests that BMAL1 contributes to the regulation of cytoskeletal factors and complexity in OPCs, similar to other glial cells (*12, 31*). Collectively, these data suggest that BMAL1 loss in OPCs could be associated with accelerated aging as both declining OPC proliferation and density and changes in morphology are linked to aging and aging-related brain disorders (*32, 33*). It should also be noted that although BMAL1 acts as a transcription factor coordinating clock-controlled genes expression (*21*), the majority of BMAL1 targets do not show rhythmic transcription patterns (*34*). While our *in vitro* findings support that OPCs are circadian dysregulated, whether BMAL1 function in OPCs is strictly circadian is an open area of investigation. Future work focused on other clock genes will distinguish non-circadian BMAL1-regulated pathways in oligodendroglia.

Previous work demonstrated that changes in astrocyte circadian clock within a demyelinating lesion signal suppression of subventricular zone BMAL1 in neural precursor cells, driving them towards the oligodendrocyte lineage (*35*). However, this work did not investigate the specific effect of BMAL1 loss on oligodendroglia in remyelination. The data presented here imply that BMAL1 controls OPC recruitment rather than proliferation of existing OPCs, a critical step for remyelination (*36*) that is deficient in MS (*37*). Decreased expression of the migratory factors *Arpc2* (*38*) and *Sox5* (*39*) and aberrant morphology following OPC-specific BMAL1 loss further supports its role in migration. BMAL1-intact oligodendroglia from older adult mice become phenotypically similar to BMAL1-disrupted oligodendroglia following demyelination (**Fig. 4G, J, L**). This suggests that disruptions in the circadian system of oligodendroglia may contribute to the limited remyelination potential associated with progressive MS and aging (*40*). OPCs from aged rats lose the ability to differentiate into oligodendrocytes, but reversing DNA damage in aged OPCs restores remyelination potential (*41*). The normal process of aging is associated with declining circadian function through changes in circadian gene expression (*42*) and BMAL1 deficiencies lead to premature aging (*43*). Whether OPCs become circadian dysregulated with age remains to be fully elucidated.

Having established that *Bmal1* expression is necessary for proper OPC lineage maintenance and myelin formation early in life, and global *Bmal1* knockouts exhibit severe circadian (*26*) and sleep phenotypes (*27*), we aimed to discern if OPC-specific BMAL1 dysregulation and consequent dysmyelination affects these systems-level processes. Mice lacking functional BMAL1 in OPCs exhibit fragmented sleep architecture during the active phase, and this fragmentation is exacerbated by sleep deprivation. Importantly, these mice do not display the gross sleep deprivation that is linked to decreased myelin sheath thickness (*44*). These findings provide a hitherto underappreciated link between myelination and healthy sleep function, a connection that may be relevant to a range of neurological diseases. In mice, sleep fragmentation increases with age (*45*) and excessive sleep fragmentation is common in age-related disorders like Alzheimer’s disease (*46*), a disorder associated with deficits in oligodendrocytes and myelin (*47*). Numerous lines of evidence support that altered circadian biology is also significantly associated with MS prevalence. Up to 60% of individuals with MS report sleep and circadian rhythm abnormalities (*48*). Our finding that sleep fragmentation is causally associated with MS prevalence further supports the conclusion that myelination may be an underappreciated regulator of sleep processes. Whether these deficits are driven by changes in BMAL1 in OPCs, OPC population integration (*8*) into neural circuitry that controls sleep, or by alterations to myelin within those circuits remains to be determined. By defining the role of the circadian system in oligodendroglial lineage cell homeostasis, we have identified new mechanistic insights into myelin and sleep regulation that may provide therapeutic targets for brain disorders.

## Acknowledgments

We would like to thank Drs. Richard Daneman and Caterina Profaci for their helpful guidance related to the assessment of the blood-brain barrier permeability.

## Funding

The U.S. Department of Defense (W81XWH-21-1-0846, EMG)

The National Multiple Sclerosis Society (PP-1907-34759, EMG)

The Weintz Family COVID-19 Research Fund (EMG)

The Department of Psychiatry and Behavioral Sciences, School of Medicine, Stanford University 2021 Innovator Grant Program (EMG)

The Brain and Behavior Research Foundation NARSAD (AW905644, EMG)

The Maternal and Child Health Research Institute Postdoctoral Fellowship (DR)

National Science Foundation Graduate Research Fellowship (DGE-1656518, CAG)

Ford Foundation Predoctoral Fellowship (CAG)

The National Cancer Institute, DHHS (PHS CA09302, LCM)

BioX Institute Fellowship (JG, MEG, JR)

The Instrumentarium Science Foundation (HMO)

NIH Shared Instrumentation Grant (1S10RR02678001; Electron Microscopy Core at Stanford University Cell Sciences Imaging Facility)

## Author contributions

Conceptualization: EMG, DR, AB, LDC. Methodology: EMG, DR, AB, LDC, SK, JG, EE, CAG, LCM, BY, SEJ, NS, HMO, SN. Investigation: EMG, DR, AB, LDC, SK, JG, EE, CAG, LCM, MEG, RS, JR, SEJ, NS. Visualization: EMG, DR, AB, SK, JG, EE. Funding acquisition: EMG. Writing: EMG, DR.

## Data and materials availability

All data are available in the main text or the supplementary materials.

## Supplementary Materials

Materials and Methods

Figs. S1 to S6

Tables S1 to S2

References (*49-59*)

