## Supplementary Material for "BMAL1 loss in oligodendroglial lineage cells dysregulates myelination and sleep"

**This PDF file includes:**

Materials and Methods
Figs. S1 to S6
Tables S1 to S2

### Materials and Methods

Animal Procedures and Mouse Models

All procedures were performed in accordance with guidelines set in place by the Stanford University Institutional Care and Use Committee. Mice were housed in group cages according to standard guidelines with *ad libitum* access to food and water under a 12::12 hr light/dark cycle. Both sexes were equally used in all experiments. *NG2::Cre* and *PDGFRα::CreER*$^{T2}$ were bred with *Bmal1*$^{fl/fl}$ mice (The Jackson Laboratory 008533, 018280, 007668) to produce *NG2::Cre;Bmal1*$^{fl/fl}$ and *PDGFRα::CreER*$^{T2}$*;Bmal1*$^{fl/fl}$ conditional constitutive and inducible knock out models, respectively. B6.129S6-*Per2*$^{tm1Jt}$/J mice (The Jackson Laboratory 006852) were bred for OPC isolation for *in vitro* bioluminescence assays. Genotyping was performed by PCR using genomic DNA extracted from ear or toe biopsies. Primers used for genotyping are listed in **Table S2**. *PDGFRα::CreER*$^{T2}$*;Bmal1*$^{fl/fl}$ mice were intraperitoneally (IP) injected with 100 mg/kg tamoxifen (Sigma T5648) for 3 consecutive days at 3 months (younger adults) or 10 months (older adults) of age. For OPC proliferation assessment, 40 mg/kg EdU (5-ethynyl-2'-deoxyuridine, Invitrogen E10187) was IP injected for 3 consecutive days before perfusions.

Lysolecithin Demyelination Model

Lysolecithin (L-α-Lysophosphatidylcholine, Sigma L4129) was diluted to 1% in sterile 1x phosphate-buffered saline (PBS), and aliquots were frozen at −20°C for single use. Six weeks after tamoxifen injections, mice were anesthetized with 3.5–5% isoflurane and placed in a stereotactic apparatus. Isoflurane was maintained at 1.5-3.0% throughout the procedure. The cranium was exposed via midline incision under aseptic conditions and a hole was drilled in the skull for injection at the following coordinates: +1.0 anterior to bregma, +1.0 lateral to midline, +1.4 dorsal to ventral. A blunt-tip 26s-gauge Hamilton syringe was used to inject 2μl of 1% lysolecithin into the cingulum of the corpus callosum of one hemisphere using a digital pump at infusion rate of 0.2 μl/min. At the completion of infusion, syringe needle remained in place for a minimum of 5 min to minimize backflow of the injection. Mice were administered bupivacaine prior to first incision, and carprofen and 0.9% sodium chloride upon completion of sutures. Mice were monitored post-operatively daily until perfused.

Telemetry Implantation and Locomotor Recording

To measure locomotor activity, 2-month old *NG2::Cre;Bmal1*$^{fl/fl}$ mice were implanted with a telemetry device (G2 E-Mitter, Mini Mitter OR, USA) in the abdominal cavity under 3% isoflurane anesthesia. Mice were allowed to recover for 3 weeks. Locomotor activity was telemetrically monitored in 12::12 light/dark (LD) cycles for 7 days followed by constant darkness (DD) for 15 days. Locomotor activity was monitored through VitalView software (Mini Mitter, OR, USA) and analyzed in 1 hour bins.

Electroencephalogram (EEG)/ Electromyogram (EMG) Surgery and Sleep Recordings

To evaluate sleep, a headstage with 4 EEG and 2 EMG electrodes on the skull was surgically implanted under 3% isoflurane anesthesia in 3.5-month old mice. Two EEG electrodes were screwed over the motor cortex (±1 mm lateral and 1 mm anterior to bregma) and the other two electrodes over the visual cortex (±3 mm lateral and 1 mm anterior to lambda). Two Teflon-coated stainless-steel wires for EMG were placed into the neck extensor muscles. At 5 months of age, mice were tethered to recording cables that connect to a low-torque slip-ring commutator (Biella Engineering, Irvine, CA, USA). After 1 week of habituation to the cable and recording environment, EEG/EMG activity was recorded for 10 days. Mice were then sleep deprived for 6 hrs, and sleep waves were recorded for 18 hrs post sleep deprivation.

EEG and EMG signals were acquired using a Glass amplifier (West Warwick, RI, USA), filtered (30 Hz Low Pass Filter for EEG; 10–100 Hz Band Pass Filter for EMG) and captured at a sampling rate of 128 Hz using a sleep recording system (Vital Recorder; Kissei Comtec Co. Ltd., Matsumoto, Japan). EEG and EMG signals were automatically scored in 10 s epochs using SleepSign software (Kissei Comtec Co. Ltd., Matsumoto, Japan). According to standard criteria, each epoch was classified as Wake, NREM, or REM sleep. The automatic scoring output was visually examined and confirmed by an experienced scorer, blind to genotype. Fast Fourier transform (FFT) was performed to analyze power spectra profiles of EEG over a 0-30 Hz with 0.5 Hz bins for the δ (0.5–4 Hz) and θ (4–9 Hz) bandwidths in a state-specific manner. A relative EEG power spectrum was calculated from EEG power densities in each frequency bin and normalized by the total bandwidth over 0-30 Hz. Epochs with artifacts and noise were excluded from the calculation.

Behavioral Tests

Motor function was tested at P35 using a CatWalk gait analysis system (Noldus, Netherlands). Mice were acclimated to handling before recording but no training was performed on the CatWalk apparatus prior to testing. Testing was performed in a dark room with red light during the light phase. Runs were considered successful when mice performed a consistent movement lasting no more than 5 s. All the analysis was performed on the CatWalk XT 9.0 software using four successful runs per mice. Swing speed was defined as the speed (cm/s) of the paw during limb swinging. It was calculated as the average of left and right forepaw swing speeds. Stride length was defined as the distance (cm) between successive placements of the same paw in the glass blade from one step cycle until the same paw touches the blade in the next cycle. It was calculated based on the X-coordinates of the center of the paw during maximum contact of the paw with the glass blade.

To specifically target frontal lobe-associated short-term memory and attention rather than long-term memory and hippocampal function in mice, a modified version of the novel object recognition task (NORT) was implemented. At 7 months of age, mice were handled for 2 minutes daily for the duration of the week leading up to the test. Mice were placed in the experimental chamber to acclimatize for 20 min the day before testing. Opaque Plexiglas experimental chambers of 61cm x 61cm x 61cm size, and a camera mounted 115 cm above the chamber were used. The tests were conducted during the first half of the animal's light phase, in a room dimly lit by standing lamps aimed away from apparatus to provide diffuse light. On the day of testing, mice were handled for 2 min before acclimatizing for 20 min in the empty chamber. Mice were returned to their home cages for 5 min before being placed in the experimental chamber with either 2 identical Lego objects (of approx. 12 cm in height) or 2 identical opaque mini bottles approximately the same width and height as the Lego-objects to counterbalance the use of each object. The position of the novel object from trial to trial was also counterbalanced. Objects were randomly assigned to avoid bias. Objects were secured in the center of the arena, 10 cm from the sidewalls and mice were placed in the center of the chamber between the 2 objects, facing perpendicular to the objects. During the training phase, mice were allowed to explore the identical objects for 5 min before being returned to their cages for 5 min while objects were cleaned with 70% ethanol. One of the 2 objects was then replaced with a new, cleaned un-identical object. During the testing phase, mice were again placed at the center of the chamber between both objects and facing perpendicular to both objects. During this phase, mice were allowed to explore for 10 min. Trials were recorded using Ethovision XT (Noldus Information Technology, Wageningen, the Netherlands) and interactions were scored by a researcher blinded to all conditions. Interactions with each object are defined as sniffing, biting and/or head within 2 cm of object, but not casually touching the object in passing or climbing. Only animals that explored the objects for a minimum of 10 s were included in the analysis.

Perfusions and Immunofluorescence

Mice were anesthetized using lethal dose of Avertin (tribromoethanol, Sigma T48402) and transcardially perfused with 15 ml PBS using a peristaltic pump (VWR 70730-062). All mice were perfused during the rest phase at Zeitgeber time (ZT)5-9 (ZT0=lights on; ZT12=lights off), with all experimental and control mice perfused at the same ZT. Brains were fixed in 4% paraformaldehyde (PFA) overnight at 4°C, and then cryoprotected in 30% sucrose. Forty µm coronal sections were taken using a sliding microtome (Epredia HM450 22-050-856). For immunohistochemistry involving EdU, sections were stained using Click-iT EdU cell proliferation kit and protocol (Invitrogen C10339) followed by incubation in 3% normal donkey serum with 0.3% Triton X-100/TBS blocking solution at room temperature for 1 hr and then incubated overnight at 4°C with primary antibodies. For all other stains, sections were incubated directly in blocking solution at room temperature for 1 hr and then incubated overnight at 4°C with primary antibodies. Goat anti-PDGFRα (1:500; R&D Systems AF1062), rabbit anti-PDGFRα (1:200, Cell Signaling Technology 5241), mouse anti-CC1 (1:100; EMD Millipore OP80), rat anti-MBP (1:200; Abcam ab7349) or rabbit anti-BMAL1 (1:500, Novus Biologicals NB100-2288) were diluted in 1% normal donkey serum in 0.3% Triton X-100 in TBS. CC1 was incubated for 7 days at 4°C as previously described (*6*). Sections were then rinsed three times in 1X TBS and incubated overnight at 4°C in secondary antibody diluted to 1:500 in 1% normal donkey serum with 0.3% Triton X-100/TBS. The secondary antibodies used include: Alexa 488 donkey anti-mouse IgG, (Jackson Immunoresearch 515-545-072), Alexa 594 donkey anti-mouse IgG, (Jackson Immunoresearch 715-585-150), Alexa 647 donkey anti-mouse IgG, (Jackson Immunoresearch 715-605-150), Alexa 488 donkey anti-rabbit IgG (Jackson Immunoresearch 711-546-152), Alexa 594 donkey anti-rabbit IgG (Jackson Immunoresearch 711-586-152), Alexa 647 donkey anti-rabbit IgG (Jackson Immunoresearch AB_2492288), Alexa 488 donkey anti-goat IgG, (Jackson Immunoresearch 705-545-147), Alexa 594 donkey anti-goat IgG (Jackson Immunoresearch 705-586-147), Alexa 647 donkey anti-goat IgG (Jackson Immunoresearch 705-606-147), Alexa 488 donkey anti-rat IgG (Jackson Immunoresearch 712-545-153), Alexa 594 goat anti-rat IgG (Jackson Immunoresearch 112-005-167). The next day, sections were rinsed 3 times in TBS, incubated with DAPI for 5 min (1:1000; Thermo Fisher Scientific 62247) and mounted with ProLong Gold mounting medium (Life Technologies P36930). Secondary-only stains were performed as negative controls.

Confocal Imaging and Quantification

Sections were imaged using a confocal microscope (Zeiss LSM900). Images were taken at 20X or 40X magnification according to the experiment and analyzed using Zen software. Z-stacks were acquired, and a maximum intensity image was generated for each frame. Cell counting was performed blinded to experimental conditions and genotype. Cells were considered co-labeled when two immunofluorescent markers co-localized in the same plane. For EdU stereology, all PDGFRα$^{+}$/EdU$^{+}$ cells were counted within the regions. The density of cells was determined by dividing the total number of PDGFRα$^{+}$/EdU$^{+}$ cells quantified for each lineage by the total volume of the imaged frames ($mm^3$). For each mouse, 6 pictures were taken in the frontal lobe corpus callosum, and all the cells were counted for each immunohistochemical marker analysis. For analysis of MBP expression, Z-stacks at 10X were analyzed using the ImageJ software (version 1.8.0). The average fluorescent intensity of pixels was quantified for the corpus callosum in 6 sections. Average values across all sections were calculated on a per animal basis.

To measure the volume of the corpus callosum, the Cavalieri Estimator function on a MBF Bioscience StereoInvestigator version 11.01.2 was used. Twelve coronal sections throughout the frontal lobe region were stained with MBP antibody and, under a 10X objective, the Cavalieri Estimator probe was used to trace the corpus callosum using published mouse anatomical guides to determine the boundaries.

For analyzing OPC morphology, semi-automated quantification was performed on Imaris for Neuroscientists Cell Imaging Software ver. 9.8. Filament Tracer Tool kit was used to automatically identify PDGFRα signal in the 3D space of 20X images. The spatial location of the cell body and processes was performed using the Autopath algorithm with the following parameters: starting points of 12.0 µm of largest diameter, and seed point of 1.62 µm of thinnest diameter. Morphological complexity was defined as the ratio between the sum of branch points and filament length by filament volume [complexity = (branch points + filament length) / filament volume].

For lysolecithin-induced demyelination studies, MBP staining was used to identify the demyelinated lesion epicenter. Two images within the lesioned area of the cingulum were taken at 20X magnification for quantification: one medial and one lateral to the demyelinated lesion epicenter. Quantification in the lesion epicenter was avoided to prevent confounds associated with needlestick injury and subsequent OPC proliferation and microglial reactivity. Cell counting was performed on Z-stacks as previously stated and analyzed using Zen software. The average cellular density of both sections was compared to the contralateral non-lesioned side.

Transmission Electron Microscopy

Mice under anesthesia with a lethal dose of Avertin were sacrificed by transcardial perfusion with 10 mL PBS followed by 10 mL Karnovsky's fixative: 2% glutaraldehyde (EMS 16000) and 4% paraformaldehyde (EMS 15710) in 0.1 M sodium cacodylate (EMS 12300), pH 7.4. A region containing premotor cortex and corpus callosum was resected from the brain and post-fixed in Karnovsky's fixative for at least 2 weeks. The effect on myelination due to OPC-specific *Bmal1* loss was measured by transmission electron microscopy of axial sections in the region of the premotor projections entering the corpus callosum at the level of the cingulum, while post-injury remyelination was measured on sagittal sections of the corpus callosum at 600 µm lateral to the lesion epicenter. First, samples were post-fixed in a solution of 2% osmium tetroxide (EMS 19100) for 5 hr at 4°C, washed 3 times with ultrafiltered water, then en bloc stained with 1% uranyl acetate overnight at room temperature and dehydrated in ethanol (30%, 50%, 70%, and 95%) for two rounds of 10 mins each at room temperature, followed by 10 minutes in 100% ethanol at 4°C, and a final step of 10 mins in propylene oxide at room temperature. Samples were infiltrated with increasing concentrations of the epoxy resin EMbed-812 (EMS 14120) at room temperature, beginning with 1 hr in 1:2 with EMbed-812:propylene oxide, followed by 1 hr in 1:1 EMbed-812:propylene oxide, then overnight in 2:1 EMbed-812:propylene oxide, then 4 hrs in pure EMbed-812. Samples were then set in TAAB capsules filled with fresh Embed-812 and left at 65°C overnight. Sections were cut at 80 nm thickness on a Leica Ultracut S (Leica, Wetzlar, Germany) and mounted on Formvar/carbon coated slot grids (EMS FCF2010-Cu) or 100 mesh Cu grids (EMS FCF100-Cu). Grids were contrast stained for 30 sec in 3.5% uranyl acetate in 50% acetone followed by staining in 0.2% lead citrate for 2 mins. Samples were imaged using a JEOL JEM-1400 TEM at 120kV and images were collected using a Gatan Orius digital camera. g-ratios were measured by dividing the axonal diameter by the diameter of the entire fiber (diameter of axon/diameter of axon + myelin sheath) at 8000X using ImageJ software blinded to genotype and condition. For each animal, 100 axons were scored. Statistical analyses were calculated using the mean g-ratio per mouse, with each mouse representing one data point. Scatter plots were performed using g-ratios as function of axon calibers of all the axons analyzed.

Blood-brain Barrier Permeability

At ZT10.5, mice were injected IP with 25 mg/kg NaFl (Sigma 46960) and, after 90 minutes, anesthetized with a lethal dose of Avertin. Blood from the right ventricle was extracted with a 23G needle and added into an EDTA-coated tube. Mice were then perfused with PBS at 8 ml/min using a peristaltic pump (VWR 70730-062) before the frontal cortex and cerebellum were micro-dissected. Blood was centrifuged to collect plasma supernatant and then diluted 1:10 in sterile PBS followed by an additional 1:1 dilution in 2% Trichloroacetic Acid (TCA). Tissue was homogenized in 1mL of PBS per 150 mg of tissue and centrifuged. Supernatants were then diluted 1:1 in 2% TCA. Both sets of samples were centrifuged, and supernatants were diluted 1:1 in borate buffer, pH 11 and fluorescence of the diluted supernatants was measured (SpectraMax iD3, excitation 480, emission 538). To ensure measurements were within detectable limits, a standard curve and blanks were used. Standards were performed with serial dilutions of lysates 0.5% TCA and 50% borate buffer in PBS with serial dilutions of sodium fluorescein dye. Blanks were 0.5% TCA and 50% borate buffer in PBS with no sodium fluorescein dye. Fluorescence ratios of brain lysates and blood samples were calculated for each mouse by taking the average of 3 technical replicates.

OPC Isolation and Culture

Mouse pups were sacrificed at P6 and brains were enzymatically and mechanically dissociated as described (*49*). OPCs were purified using immunopanning through 2 negative selection plates for microglia and endothelial cells with BSL1 (Vector Laboratories L1100) and one positive selection plate with an anti-PDGFRα antibody (rat anti-mouse CD140A, BD Pharmingen 558774) as described (*49*). OPCs were seeded in Poly-D-lysine coated plates (Sigma P6407) and incubated at 37°C, 10% CO2 in DMEM-Sato base growth medium composed of Dulbecco's Modified Eagle's Medium (Invitrogen 11960-044), 100 U/mL penicillin and 100 µg/mL streptomycin (Gibco/Life Technologies 15140-122), 2 mM Glutamine (Invitrogen 25030-081), 1 mM Sodium pyruvate (Invitrogen 11360-070), 5 µg/mL Insulin (Sigma-Aldrich I6634), 5 µg/mL N-Acetyl-L-cysteine (Sigma-Aldrich A8199), 1x Trace Elements B (Corning 25022CI), 10 ng/mL d-Biotin (Sigma-Aldrich B4639), 100 µg/mL BSA (Sigma-Aldrich A4161), 100 µg/mL Transferrin (Sigma-Aldrich T1147), 16 µg/mL Putrescine (Sigma-Aldrich P5780), 60 ng/mL Progesterone (Sigma-Aldrich P8783), 40 ng/mL Sodium selenite (Sigma-Aldrich S5261), 4.2 µg/mL Forskolin (Sigma-Aldrich F6886) and 10 ng/mL CNTF (Peprotech 450-13). To promote OPC proliferation, cells were maintained in 10 ng/mL PDGF-AA (Peprotech 100-13A) and 1 ng/mL NT-3 (Peprotech 450-03). To promote OPC differentiation, cells were maintained in 1x B-27 (Invitrogen 17504-044) and 40 ng/mL T3 (Sigma-Aldrich T6397). Half of the media was replaced with fresh media every 2 days.

OPC Immunofluorescence

OPCs were seeded in ethanol-washed glass coverslips coated with Poly-D-lysine (Sigma P6407) in a density of 5000 cells/well in proliferation or differentiation media. For analyzing OPC proliferation, EdU was added to the media for 5 hrs and cells were then fixed in 4% PFA for 15 min at room temperature. For EdU staining, sections were permeabilized with 0.1% Triton X-100 in PBS for 3 min at room temperature and stained using Click-iT EdU cell proliferation kit and protocol (Invitrogen C10339) followed by an incubation with 3% BSA in PBS for 20 min at room temperature. For differentiation assays, after 3 or 6 days of OPCs incubation with differentiation factors, cells were fixed in 4% PFA for 15 min at room temperature, permeabilized with 0.1% Triton X-100 in PBS for 3 min at room temperature followed by an incubation with 3% BSA in PBS for 20 min at room temperature. Rat anti-MBP (1:500; Abcam ab7349), rabbit anti-BMAL1 (1:500, Novus Biologicals NB100-2288), goat anti-PDGFRα (1:500; R&D Systems AF1062) or rabbit anti-NG2 (1:500, Millipore AB5320) were diluted in 3% BSA in PBS and incubated overnight at 4°C. The next day, coverslips were rinsed 3 times in PBS and incubated in secondary antibody solution in 3% BSA for 2 hr at room temperature. The following secondary antibodies were used at 1:500: Alexa 594 goat anti-rat IgG (Jackson Immunoresearch 112-005-167), Alexa 488 donkey anti-rabbit IgG (Jackson Immunoresearch 711-546-152), Alexa 488 donkey anti-goat IgG (Jackson Immunoresearch 705-545-147), Alexa 594 donkey anti-rabbit IgG (Jackson Immunoresearch 711-586-152). Coverslips were rinsed 3 times in PBS, incubated with DAPI for 10 min (1:1000; Thermo Fisher Scientific 62247) and mounted with ProLong Gold mounting medium (Life Technologies P36930). For differentiation assays, Phalloidin (1:143 of 66 μM stock, Invitrogen A12379) was also added to the DAPI solution to stain F-actin to distinguish the differentiation stage as stated in (*50*). Secondary-only stains were performed as negative controls. Confocal Imaging and quantification were performed as previously stated for *in vivo* experiments on Z-stacks from 3x3 tiles taken at 10X.

Western Blots

Isolated OPCs were lysed with RIPA buffer containing protease inhibitors. Lysates were then centrifuged at 13,000g for 15 min at 4°C and total protein was quantified (Biorad 5000113 & 5000114). All samples were normalized by protein concentration, mixed with 1X Bolt DTT Reducing Agent (Invitrogen #B0009) and 1X Bolt LDS sample buffer (Invitrogen B0007), incubated at 70°F for 10 min, and loaded onto Bolt 4-12% Bis-Tris Plus (Invitrogen NW04120). Protein was semi-dry transferred at 20-25V onto nitrocellulose membranes (Invitrogen IB23002) and blocked with 5% bovine serum albumin (BSA Sigma #A3294) in TBST for 1h. Direct-blot™ HRP rat anti-GAPDH (1:1000, BioLegend 607903) and rabbit anti-BMAL1 (1:2000, Novus Biologicals NB100-2288) antibodies were diluted in 1% BSA/TBST and membranes incubated overnight at 4°C. The following day, membranes were rinsed 3 times in TBST and incubated with a secondary goat anti-rabbit IgG antibody conjugated to horseradish peroxidase (1:5000, VWR 10150732) for 1hr. Proteins were visualized using WesternBright Sirius (Advansta K-12043-C20) and quantified using Image Studio Lite software (LI-COR).

Luminescence Assays

To test OPC circadian synchronization *in vitro*, immunopan-isolated OPCs from B6.129S6-*Per2*$^{tm1Jt}$/J P6 mouse brains were seeded in 60000 cells/well in 96 well opaque plates. OPCs were treated with 100 nM dexamethasone (Sigma-Aldrich D4902) for 1 hr followed by a complete media change or synchronized using a media change containing 10 μM forskolin (Sigma-Aldrich F6886). At circadian time (CT)4, 8, 12, 16, 20 and 24 after *in vitro* circadian synchronization, 1 μM beetle luciferin potassium salt (Promega E1602) was added, and bioluminescence was recorded after 20 min in a SpectraMax iD3 plate reader.

RT-qPCRs

Clock gene expression was measured on primary cultures of OPCs isolated from P6 mouse brains and circadian synchronized *in vitro* using dexamethasone 100nM for 1 hr. Samples were collected at CT4, 8, 12, 16, 20 and 24 after *in vitro* synchronization, lysed with 500 µL QIAzol Lysis Reagent (Qiagen 79306) and stored at -80°C until RNA extraction. For expression of genes associated to cytoskeleton, cell cycle or OPC proliferation and differentiation regulation, OPCs from P6 mouse brains were isolated at ZT6 using immunopanning and one million cells were immediately lysed using QIAzol Lysis Reagent, as previously stated. RNA extraction was performed following manufacturer's instructions and RNA purity and concentration were assessed using a Nanodrop and checking 260/280 and 260/230 nm ratios. 600 ng of RNA were used for first-strand cDNA synthesis using Oligo-dT primers and SuperScript™ III First-Strand Synthesis System (Invitrogen 18080051). Samples diluted in 1:8 were run in triplicate in an Applied Biosystems QuantStudio™ 6 Flex Real-Time PCR System machine using Power Up SYBR Green Master Mix (Applied Biosystems A25776) following supplier's protocol and a cycling program of 2 min at 50°C followed by 2 min at 95°C and 40 cycles of 95°C for 15 sec and 60°C for 1 min. Primers used are listed in **Table S2**. Melting curves were analyzed to confirm specificity of the PCR product. Relative quantification against *Gapdh* reference gene expression was done using the 2-ΔΔCT method.

Mendelian randomization (MR) analyses

MR analyses were performed using genetic variants previously reported to be associated with Multiple Sclerosis (*30*), and instruments from the mean number of sleep episodes per sleep period (a proxy for sleep fragmentation) (*51*), which was measured through wrist-worn accelerometer devices. All variants were first corrected for Winner's Curse (*52*, *53*) using a previously described method (*54*), as Winner's Curse leads to an overestimation of the effect of the genetic instruments on the exposure and, in MR, can bias the causal effect towards the null (*55*). Of the 21 lead variants available for sleep fragmentation, two were unavailable in the MS summary statistics and further eight were excluded due to being weak instruments after Winner's Curse correction (**Table S1**). Furthermore, we compared the effect with winner's curse correction to the full set of variants without correction.

Five MR approaches were used: inverse-variance weighted (IVW), Weighted median, Weighted mode, Simple mode and MR with Egger regression (MR Egger) using TwoSampleMR package version 0.5.6 with R version 4.1.3. IVW is the most statistically powerful of the three methods but assumes that all included instruments are non-pleiotropic (*56*). The weighted median (WM) approach provides a causal estimate that is consistent even if up to half of the variants are invalid instruments (due to pleiotropy), with the caveat of a loss of statistical power (*57*). The MR Egger approach loosens the requirement that all genetic variants satisfy the exclusion-restriction principal. In the context of MR, this allows a subset of variants to be pleiotropic and so adjusts the causal estimate for the estimated overall directional pleiotropy (represented by a constant term in the linear regression equation) (*58*). Power calculations were performed using the mRnd tool (*59*), available at the website https://shiny.cnsgenomics.com/mRnd/, using the Binary Outcome calculator with the total sample size set as 115,803 (sample size of the Multiple Sclerosis consortium), alpha as 0.05 (default), the proportion of cases in the study as ˜0.69 (= 47429/68374), the true odds ratio as 1.17 (the Weighted median causal effect size estimate) and the proportion of variance explained as ~0.004674 (the sum of the variance explained by the 11 variants used in the MR analysis resulting 38% power to detect association with this alpha level; **Table S1**).

Data Availability

The Multiple Sclerosis summary statistics are publicly available at through TwoSampleMR package. To access individual-level data from the UK Biobank studies, researchers must apply for access through the UK Biobank portal at https://www.ukbiobank.ac.uk/enable-your-research/apply-for-access.

Statistical Analysis

All statistical analyses were conducted using GraphPad Prism statistical software, including Tests of Normality. Group mean differences between 2 genotypes for all immunofluorescence and Western blot analysis, morphological complexity, g-ratios, corpus callosum volume, RT-qPCR assays at ZT6, CatWalk behavioral tests (swing speed and stride length), circadian period length, and locomotor activity were assessed using unpaired, two-tailed Student's t-tests. Group mean differences between genotypes in proliferation and differentiation *in vitro* assays were assessed using nested t-test to account for different experimental replicates. Group mean differences between 2 genotypes and different circadian times for RT-qPCR of clock genes were compared with 2-way ANOVA with Holm-Sidak post-hoc tests to further examine pairwise differences. Group mean differences between 2 genotypes and 2 conditions for NORT behavioral tests, sleep recordings, analysis of mean differences between the ipsilateral and contralateral sides within the same animal in the lysolecithin model, and blood-brain barrier permeability analysis were assessed with 2-way ANOVA with Holm-Sidak and Tukey post-hoc tests to further examine pairwise differences. A level of $P < 0.05$ was used to designate significant differences.

**Fig. S1.**

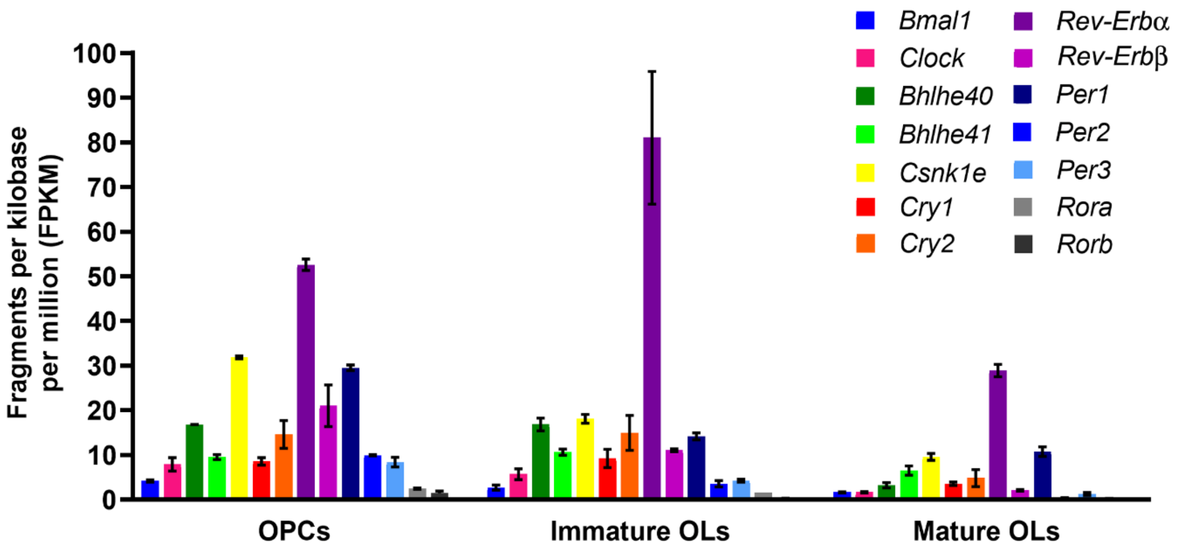


**Fig. S1.** Circadian gene expression as determined by RNA-sequencing in oligodendrocyte lineage cells isolated from P16 mice throughout the circadian day (*16*). The graph represents a composite of gene expression data from multiple circadian phases, confirming the overall presence of all core circadian clock genes throughout the lineage irrespective of phase information.

**Fig. S2.**

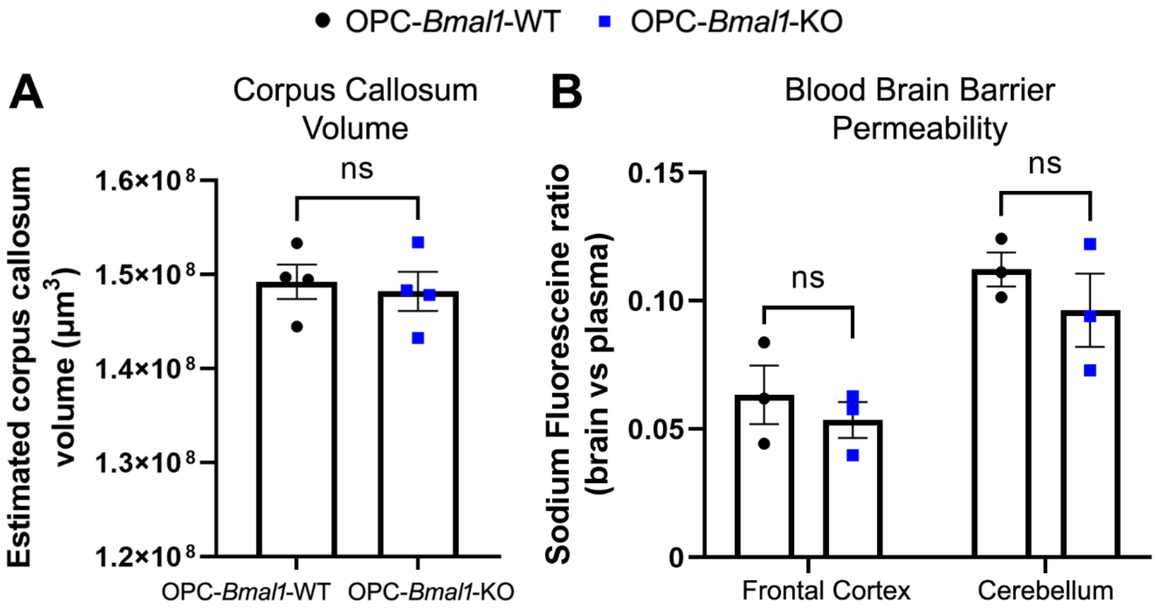


**Fig. S2. Knock out of *Bmal1* in OPCs is not associated with changes in corpus callosum volume or blood-brain barrier permeability. (A)** Constitutive knock out of *Bmal1* in OPCs (OPC-*Bmal1*-KO) does not affect corpus callosum volume compared to their control littermates (OPC-*Bmal1*-WT) (n = 4 per group). **(B)** OPC-*Bmal1*-KO mice have similar blood-brain barrier permeability compared to OPC-*Bmal1*-WT mice as assessed by measuring concentrations of sodium fluoresceine in the frontal cortex or cerebellum following systemic administration, indicating intact integrity of the blood-brain barrier at baseline (n = 3 per group). Data shown as mean +/- S.E.M. n.s. p>0.05.

**Fig. S3.**

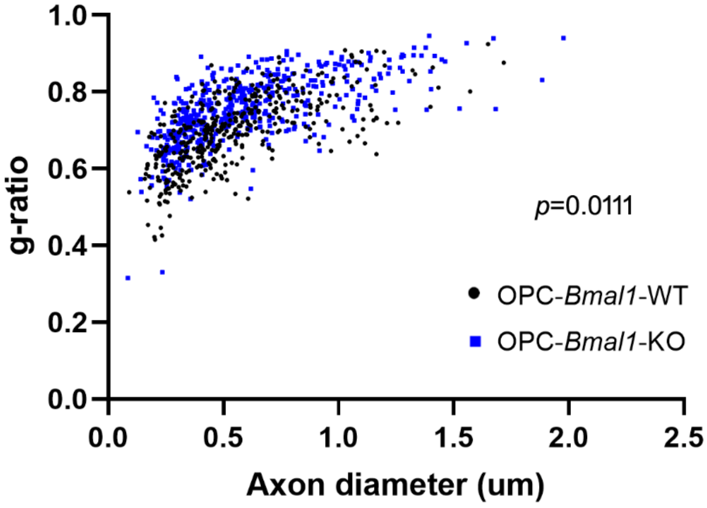


**Fig. S3. Knock out of *Bmal1* in OPCs leads to deficits in myelination.** Embryonic loss of *Bmal1* in OPCs (OPC-*Bmal1*-KO) results in decreased myelin sheath thickness of axons in the corpus callosum at P21 compared to controls (OPC-*Bmal1*-WT). Scatterplot of *g*-ratios as a function of axon caliber of all axons measured (larger g-ratio = thinner myelin). Each point represents a single axon. n = 4-5 mice per group.

**Fig. S4.**

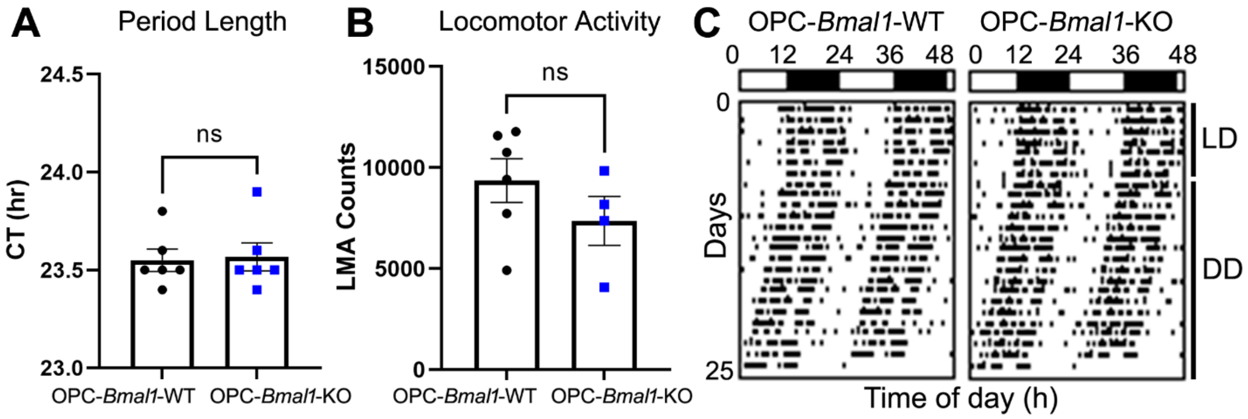


**Fig. S4. Lack of *Bmal1* in OPCs does not affect global circadian periodicity or locomotor activity. (A-B)** Knock out of *Bmal1* in OPCs (OPC-*Bmal1*-KO) does not alter circadian period length (tau) **(A)** or locomotor activity (LMA) **(B)** when mice are placed in constant darkness compared to control siblings (OPC-*Bmal1*-WT) at 3 months of age (n= 6 per group). **(C)** Representative actogram for locomotor activity in OPC-*Bmal1*-WT and OPC-*Bmal1*-KO mice in 7 days of light/dark cycle (LD) and 18 days of constant darkness (DD). Data shown as mean +/- S.E.M. n.s. p>0.05.

**Fig. S5.**

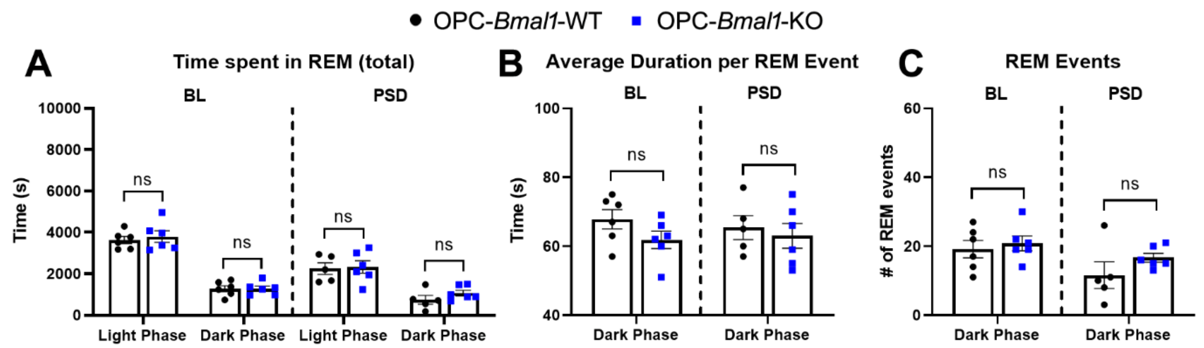


**Fig. S5. Loss of *Bmal1* in OPCs does not affect REM sleep. (A)** OPC-*Bmal1*-WT and OPC-*Bmal1*-KO spend equivalent amounts of time in REM during both the light/sleep and dark/active phases at baseline (BL) and after 6-hr sleep deprivation (PSD). **(B)** The average duration per REM event and **(C)** the total number of REM events do not differ between the genotypes. n= 5-6 per group. Data shown as mean +/- S.E.M. n.s. p>0.05.

**Fig. S6.**

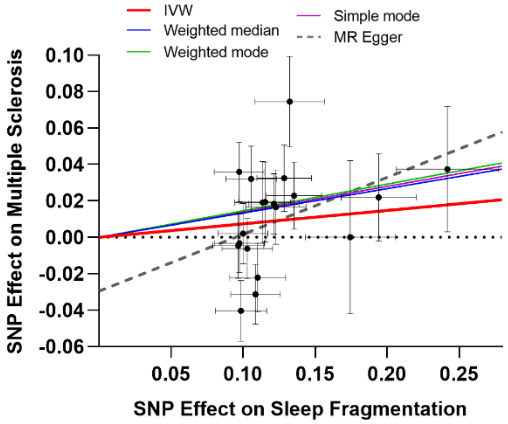


**MR analysis of association between sleep fragmentation and multiple sclerosis**

| Method | N of SNPs | Beta | SE | *P* value |
|---|---|---|---|---|
| Inverse variance weighted | 19 | 0.074 | 0.050 | 0.143 |
| Weighted median | 19 | 0.134 | 0.055 | 0.016 |
| Weighted mode | 19 | 0.146 | 0.085 | 0.104 |
| Simple mode | 19 | 0.140 | 0.100 | 0.178 |
| MR Egger | 19 | 0.312 | 0.233 | 0.199 |
| MR Egger Intercept | 19 | -0.029 | 0.282 | 0.311 |

**Fig. S6. Sleep fragmentation is consistently causally associated with multiple sclerosis without excluding invalid variants.** Scatterplot of the mendelian randomization (MR) analysis of association between sleep fragmentation and multiple sclerosis using 19 variants without correcting for winner's curse. The slopes of the regression lines represent the causal association tested using Inverse variance weighted (IVW), Weighted median, Weighted mode, Simple mode, and MR Egger statistical tests. Corresponding table of the MR analysis performed indicates causal association between sleep fragmentation and multiple sclerosis risk without pleiotropic effect. Beta, β coefficient; SE, Standard Error. Data shown as mean +/- S.E.

**Table S1.**

**Table S1. Summary statistics for the Mendelian randomization analysis performed for each genetic variant.** Chr, chromosome; Beta, beta coefficient; SE, standard error; logOR, log off ratios; IVW, Inverse variance weighted.

### Table S1

| | | | | | | Exposure (N Sleep Episodes) | | | | | | Outcome (Multiple Sclerosis) | | | Included in analysis |
|---|---|---|---|---|---|---|---|---|---|---|---|---|---|---|---|
| Variant | Chr | Pos_b37 | EA | OA | EAF | Beta | SE | Beta (InvNorm) | SE (InvNorm) | P | PropVarExpl | LogOR | SE | P | IVW and Weighted Median |
| rs12714404 | 2 | 282462 | T | G | 0.2832 | 0.1353 | 0.019 | 0.037 | 0.0052 | 1.70E-12 | 0.000599 | 0.0228 | 0.0185 | 0.2179 | Yes |
| rs310727 | 3 | 4336589 | T | C | 0.4748 | 0.0972 | 0.017 | 0.026 | 0.0047 | 2.00E-08 | 0.000274 | 0.0359 | 0.0163 | 0.0277 | No |
| rs55754932 | 3 | 87847754 | C | A | 0.2837 | 0.1369 | 0.019 | 0.037 | 0.0052 | 2.00E-12 | 0.000482 | - | - | - | No |
| rs9864672 | 3 | 1.37E+08 | T | C | 0.5219 | 0.1086 | 0.017 | 0.029 | 0.0047 | 1.70E-10 | 0.000357 | -0.0312 | 0.0164 | 0.5219 | Yes |
| rs4974697 | 4 | 2473092 | T | A | 0.3901 | 0.0965 | 0.018 | 0.026 | 0.0048 | 5.90E-08 | 0.000255 | -0.0046 | 0.0179 | 0.7970 | No |
| rs7377083 | 4 | 1.03E+08 | A | C | 0.4296 | 0.1055 | 0.017 | 0.029 | 0.0047 | 1.90E-09 | 0.000342 | 0.0320 | 0.0181 | 0.0774 | Yes |
| rs749100 | 5 | 63307862 | A | G | 0.5816 | 0.1214 | 0.017 | 0.033 | 0.0047 | 5.30E-12 | 0.000423 | 0.0183 | 0.0166 | 0.2682 | Yes |
| rs9341399 | 6 | 73773644 | C | T | 0.9364 | 0.2418 | 0.035 | 0.066 | 0.0096 | 1.00E-11 | 0.000456 | 0.0374 | 0.0344 | 0.2765 | Yes |
| rs1889978 | 6 | 1.25E+08 | C | T | 0.4855 | 0.0999 | 0.017 | 0.027 | 0.0047 | 5.00E-09 | 0.000352 | 0.0021 | 0.0166 | 0.8993 | Yes |
| rs2141277 | 7 | 39099178 | A | G | 0.4775 | 0.0974 | 0.017 | 0.026 | 0.0047 | 9.90E-09 | 0.000347 | -0.0032 | 0.0161 | 0.8425 | No |
| rs10233848 | 7 | 1.03E+08 | G | A | 0.2933 | 0.1285 | 0.019 | 0.035 | 0.0051 | 1.40E-11 | 0.000497 | 0.0325 | 0.0181 | 0.0725 | Yes |
| rs1124116 | 10 | 99371147 | A | G | 0.7295 | 0.1136 | 0.019 | 0.031 | 0.0052 | 3.20E-09 | 0.000344 | 0.0192 | 0.0224 | 0.3916 | Yes |
| rs4755731 | 11 | 43685168 | G | A | 0.4309 | 0.1028 | 0.017 | 0.028 | 0.0047 | 3.20E-09 | 0.000391 | -0.0062 | 0.0165 | 0.7065 | Yes |
| rs3751837 | 16 | 3583173 | C | T | 0.7805 | 0.1227 | 0.021 | 0.033 | 0.0057 | 4.10E-09 | 0.000431 | 0.0166 | 0.0204 | 0.4160 | Yes |
| rs8045740 | 16 | 20262776 | G | T | 0.8676 | 0.194 | 0.026 | 0.052 | 0.0070 | 5.80E-14 | 0.000636 | 0.0220 | 0.0241 | 0.3609 | No |
| rs11078917 | 17 | 37746359 | A | C | 0.2791 | 0.1064 | 0.019 | 0.029 | 0.0052 | 2.40E-08 | 0.000460 | - | - | - | No |
| rs11082030 | 18 | 35501739 | T | C | 0.7249 | 0.11 | 0.019 | 0.030 | 0.0052 | 8.40E-09 | 0.000383 | -0.0221 | 0.0188 | 0.2385 | No |
| rs8098424 | 18 | 52458218 | G | A | 0.6195 | 0.0985 | 0.018 | 0.027 | 0.0048 | 2.20E-08 | 0.000372 | -0.0403 | 0.0167 | 0.0156 | No |
| rs76753486 | 19 | 42684264 | T | C | 0.0836 | 0.1744 | 0.031 | 0.047 | 0.0085 | 2.00E-08 | 0.000339 | 0.0001 | 0.0420 | 0.9981 | No |
| rs429358 | 19 | 45411941 | T | C | 0.848 | 0.1323 | 0.024 | 0.036 | 0.0065 | 3.20E-08 | 0.000378 | 0.0745 | 0.0248 | 0.0026 | No |
| rs12479469 | 20 | 61145196 | A | G | 0.3419 | 0.1153 | 0.019 | 0.031 | 0.0050 | 5.60E-10 | 0.000482 | 0.0194 | 0.0219 | 0.3744 | Yes |

**Table S2.**

| Genotyping Primers | Sequence (5'-3') |
|---|---|
| *Bmal1-Flox-F* | GGGGAGGGTGAGAAAACAGA |
| *Bmal1-Flox-R* | TCCCTTGAAATTTTCTGACCAAC |
| *Cre-F* | GATCTCCGGTATTGAAACTCCAGC |
| *Cre-R* | GCTAAACATGCTTCATCGTCGG |
| **RT-qPCR Primer** | **Sequence (5'-3')** |
| *Gapdh-F* | AGGTCGGTGTGAACGGATTTG |
| *Gapdh-R* | TGTAGACCATGTAGTTGAGGTCA |
| *Bmal1-F* | ACATCACAAGTACGCCTCCC |
| *Bmal1-R* | TGCTGCCTCATCGTTACTGG |
| *Per2-F* | ATGTGCAGCCAGAACTTCCT |
| *Per2-R* | AGAGTGTGGTGTCCCTCACC |
| *Rev-Erba-F* | ACAGTGATGTTCCTGAGCCG |
| *Rev-Erba-R* | TTGGTGAAGCGGGAAGTCTC |
| *Actb-F* | TACCACCATGTACCCAGGC |
| *Actb-R* | CTCAGGAGGAGCAATGATCTTGAT |
| *Cttn-F* | ATTGAGGCCGTAACCAGCAA |
| *Cttn-R* | TCTTTCTTTGGCCATCCGCT |
| *Arpc2-F* | ATCAGGCTGGCATGTTGAAG |
| *Arpc2-R* | ACACCTTTCCGATGACCACA |
| *Sox5-F* | AATGAGCCAGAAGACACTCCCAGT |
| *Sox5-R* | AGGAGGGAACACGGGAATCATCAA |
| *Sox6-F* | TGGGCAAAGGACGAAAGGAG |
| *Sox6-R* | GCCTGTTCTTCATAGTAAGGTTGCT |
| *Id2-F* | TCCCTTCTGAGCTTATGTCGA |
| *Id2-R2* | TCAACGTGTTCTCCTGGTGA |
| *Sox10-F* | GACCAGTACCCTCACCTCCA |
| *Sox10-R* | CGCTTGTCACTTTCGTTCAG |
| *Olig2-F* | CAGCGAGCACCTCAAATCTA |
| *Olig2-R* | AGGAGGTGCTGGAGGAAGAT |
| *Myrf-F* | CCTGTGTCCGTGGTACTGTG |
| *Myrf-R* | TCACACAGGCGGTAGAAGTG |
| *c-Myc-F* | TGAGGAGACACCGCCCAC |
| *c-Myc-R* | CAACATCGATTTCTTCCTCATCTTC |
| *Cdk1-F* | GCCAGAGCGTTTGGAATACC |
| *Cdk1-R* | CAGATGTCAACCGGAGTGGAGTA |
| *Tp53-F* | ACGCTTCTCCGAAGACTGG |
| *Tp53-R* | AGGGAGCTCGAGGCTGATA |

**Table S2. Primers used for genotyping mice and for RT-qPCRs.**
